# The evolution of separate sexes in waterhemp is associated with surprising chromosomal diversity and complexity

**DOI:** 10.1101/2024.09.17.613486

**Authors:** Julia M. Kreiner, Jacob S. Montgomery, Marco Todesco, Natalia Bercovich, Yunchen Gong, Cassandra Elphinstone, Patrick J. Tranel, Loren H. Rieseberg, Stephen I. Wright

**Affiliations:** Department of Ecology & Evolution, University of Chicago; Department of Botany, University of British Columbia; Center for the Analysis of Genome Evolution and Function, University of Toronto; Department of Crop Sciences, University of Illinois Urbana-Champaign; Department of Ecology & Evolutionary Biology, University of Toronto

## Abstract

The evolution of separate sexes is hypothesized to occur through distinct pathways involving few large-effect or many small-effect alleles. However, we lack empirical evidence for how these different genetic architectures shape the transition from quantitative variation in sex expression to distinct male and female phenotypes. To explore these processes, we leveraged the recent transition of *Amaranthus tuberculatus* to dioecy within a predominantly monoecious genus, along with a sex-phenotyped population genomic dataset, and six newly generated chromosome-level haplotype phased assemblies. We identify a ∼3 Mb region strongly associated with sex through complementary SNP genotype and sequence-depth based analyses. Comparative genomics of these proto-sex chromosomes within the species and across the Amaranthus genus demonstrates remarkable variability in their structure and genic content, including numerous polymorphic inversions. No such inversion underlies the extended linkage we observe associated with sex determination. Instead, we identify a complex presence/absence polymorphism reflecting substantial Y-haplotype variation—structured by ancestry, geography, and habitat— but only partially explaining phenotyped sex. Just over 10% of sexed individuals show phenotype-genotype mismatch in the sex-linked region, and along with observation of leakiness in the phenotypic expression of sex, suggest additional modifiers of sex and dynamic gene content within and between the proto-X and Y. Together, this work reveals a complex genetic architecture of sex determination in *A. tuberculatus* characterized by the maintenance of substantial haplotype diversity, and variation in the expression of sex.

## Introduction

Dioecy is thought to primarily evolve from cosexuality through two distinct pathways (gynodioecy and monoecy-paradioecy) that have contrasting predictions about the importance of small versus large effect mutations [1]. In the gynodioecy pathway, the intermediate stage is typically characterized by a single large-effect male sterility mutation that invades a cosexual population and transitions it to polymorphic for females and hermaphrodites. This is eventually resolved to dioecy by the invasion of a female sterility mutation tightly linked to a restorer of male fertility [2]. Recombination suppression is thought to evolve to prevent fully sterile or hermaphroditic genotypes, and may be further extended due to sexually antagonistic selection that assembles male and/or female beneficial alleles on alternate haplotypes [3]. This scenario describes the evolution of an XY system, but explains a ZW system if sexes are switched. The lack of recombination in the sex-linked region is thought to drive the degeneration of the non-recombining Y/W chromosomes, although contemporary work shows extensive variation in size and the degree of degeneration, which is not a simple function of age [4] [5–9]. Nonetheless, young sex determining systems in the early stages of this process more often tend to demonstrate incomplete recombination suppression and leaky sex determination, despite multiple sexually antagonistic polymorphisms having evolved [10,11].

In dioecious species that evolve through the monoecy-paradioecy pathway, selection is thought to act on quantitative variation for sex allocation. Populations transition from individuals which are equally invested in male and female function to those alternately fixed for one or the other through gradual disruptive selection on the proportion of male and female flowers produced [12–15]. Tests of this route to separate sexes have been limited in part due to the relative infrequency of dioecious species in predominantly monoecious plant genera and families (but see *Sagittaria* [1]*, Urtica* [16], and *Populus* [17]). However, a notable example of this pathway is *Mercurialis annua*, where recent work [18] has resolved that inbreeding avoidance drives disruptive selection on phenotypic sex allocation and the evolution of dioecy. Accompanying models suggest that while polygenic variation for sex allocation may be initially required for quantitative sex expression, resolution of the monoecy-paradioecy pathway can result in the emergence of single-locus sex determination [14], which in part, could result from shifts in the recombination landscape.

While sexual intermediacy (e.g. subdioecy [19]) occurs during the evolution of dioecy itself, selection subsequent to the evolution of separate sexes may also act to maintain variation in sex expression [13,20–25]. Dioecious species are prone to this “leakiness”, even ones with seemingly long-evolved sex-determining regions [26]. While little is known about the genetic basis of leakiness, it is likely that as for the paradioecy pathway, multiple genomic loci [27] and/or expression modifiers [22] govern this phenotypic variation for sex, alongside environmental factors. Therefore, recent transitions to dioecy from monoecy in particular, provide the opportunity to investigate the genetic architecture of sex determination, with implications for understanding the timescale and evolution of recombination suppression, sex allocation, and sex expression.

The genus *Amaranthus* (Amaranthaceae) contains about 70 species, of which the vast majority are monoecious. There have likely been multiple relatively recent transitions to dioecy, as two to three dioecious clades are spread across two out of the three subgenera (*Amaranthus* and *Acnida*, but not *Albersia*) [28–31]. All dioecious species in this genus are thought to result from a male heterogametic (XY) system, with homomorphic (lack of apparent morphological differences) sex chromosomes [32,33]. The genus is economically important, as it contains both domesticated pseudograin crops and invasive agricultural weeds, the latter encompassing the focal species for this study, the wind-pollinated annual, *A. tuberculatus* (common waterhemp). *Amaranthus tuberculatus* can hybridize with monoecious relatives at relatively high rates under field conditions, highlighting the recent evolution of dioecy in the clade [32,34,35]. Interspecific hybrids between *A. tuberculatus* and monoecious species are also dioecious, suggesting dominance of dioecy in the genus [36]. Understanding the genetic basis of sex is particularly important in the *Amaranthus* genus, with implications for optimizing artificial selection and improving crop breeding efficiency (e.g. through the development of a male-sterile breeding system in domesticated species [37,38]), and informing evolutionary management approaches such as gene drive or gene silencing to suppress females and accelerate local extirpation of invasive weed populations [36,39].

Work on the causes and consequences of sex in *A. tuberculatus* illustrates sexual dimorphism, apparent at both the phenotypic and genomic level, despite the likely recent origin of separate sexes. Phenotyped males and females are differentiated in key life history traits, with males being faster growing and earlier flowering than females, and with sex differences in genome size and content of particular repeat classes [40,41]. Furthermore, levels of sexual dimorphism between males and females vary by habitat type (whether plants were collected from natural or agricultural habitats) [41], suggesting that sexually antagonistic selection is heterogeneous across environments. Recent genomic work confirmed a male heterogametic system, with a cumulative 4.6 Mb of male-specific sequence identified through sex-specific RAD-tags [36], with loci spread across multiple contigs. Following this work, Raiyemo et al. [42] produced a phased reference genome identifying chromosome 1 as enriched for sex linked SNPs, although their assemblies lacked a contiguous sex-linked region based on genome wide association (GWA) and F_ST_ analyses.

Here, we generate six new haplotype-phased assemblies from across the geographic range, and analyze them along with considerable resequencing resources to investigate the diversity and evolution of proto-sex chromosomes in *A. tuberculatus*. We first implement population genomic approaches to resolve the location of the sex-linked region along the same chromosome identified by [42]. Leveraging our multiple assemblies and comparative genomic approaches, we show sex determination in *A. tuberculatus* to be linked to a polymorphic region ∼3 Mb long, not found within an inversion, but rather differentiated by presence/absence across the sexes. The lack of complete 1:1 mapping of phenotype onto any genotype across the genome suggests that additional modifiers may be contributing to variation in sex expression in the system, such as an individual we observe to possess both male and female flowers. We find that the substantial diversity in sex-linked haplotypes are structured by ancestry, geography, and habitat, which may reflect the on-going generation of presence/absence variation or on-going gene exchange between the proto-X and -Y. Taken together, our results reveal the diversity underlying sex in this recently evolved proto-sex chromosome system.

## Results

### Characterization of the copy number polymorphic sex-linked region

To identify the location of sex-linked region(s) in our newly produced reference genomes, we performed a GWA using 186 sex-phenotyped individuals (96 female, 92 male) from paired environmental collections [40] that we mapped to one randomly chosen haplotype (haplotype 2) of the highest coverage (70X) assembly. This genome came from a male plant from Walpole, Ontario, Canada, a population admixed for ancestry between the two varieties of *A. tuberculatus*, var. *rudis* and var. *tuberculatus*. After mapping, calling, and filtering SNPs (see methods), a GWA identified Chromosome 1, the largest across the genome, as highly enriched for sex-linked alleles, consistent with recent work ([42]; **Sup Figure 1, Sup Figure 2)**.

**Figure 1.**
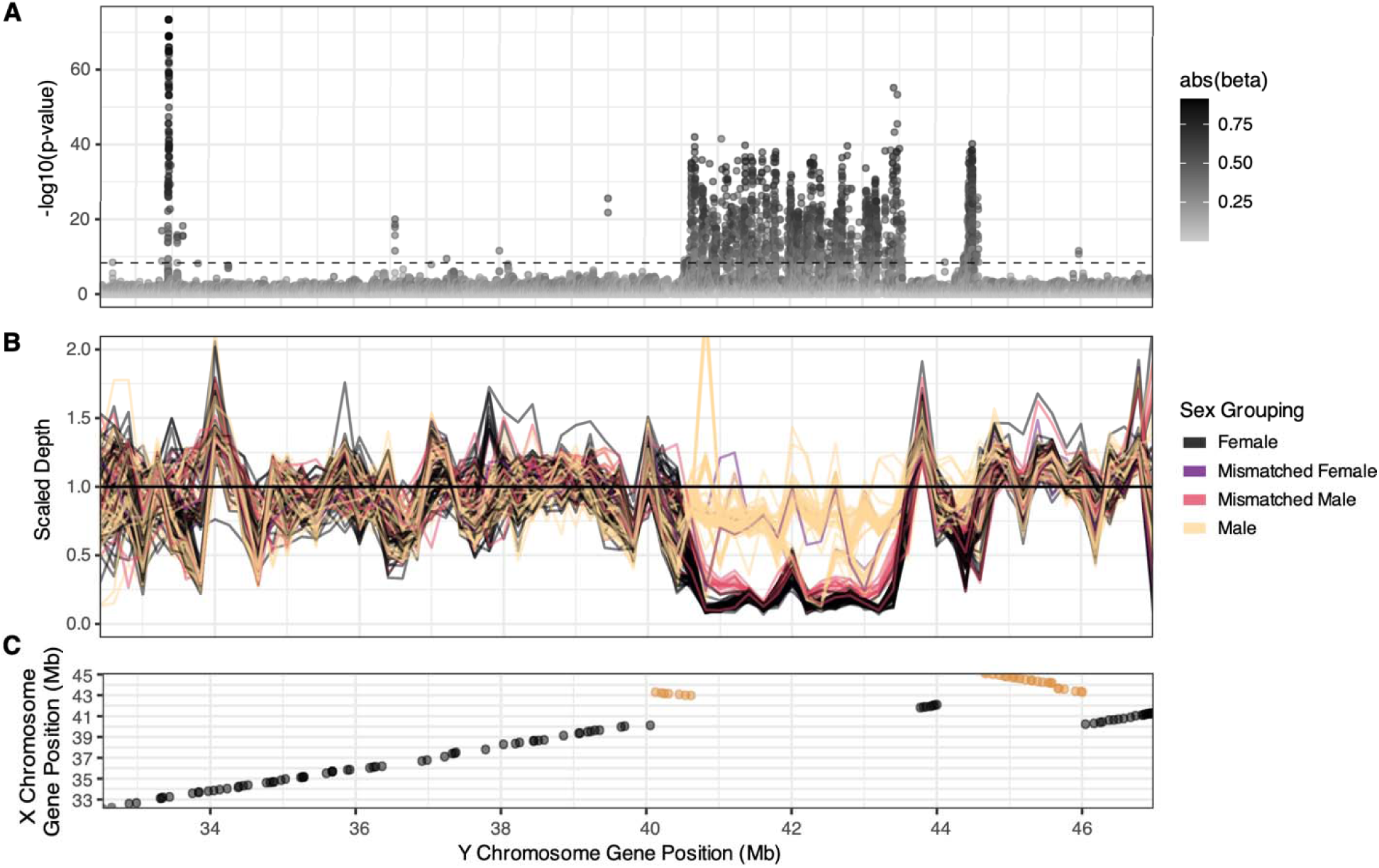
The sex-linked region (SLR) in *A. tuberculatus* on Chromosome 1. **A**) A GWA resolved strongly sex-linked alleles along a 3 Mb stretch on haplotype 2 of Chromosome 1, between 40.65-43.5[44.5] Mb. Color of points depicts the absolute allelic effect size from a mixed model. The horizontal dashed bar represents the Bonferroni p=0.05 threshold. **B**) Scaled depth individual profiles across Chromosome 1 (reflecting 200 kb moving averages), illustrating sex-based differentiation in copy number, fine-scale variation within sexes, and genotype-phenotype mismatch. **C**) Genes falling within the SLR on the Y (haplotype 2 assembly) lack syntenic orthologous genes on the X (haplotype 1 assembly). Orange represents inverted orthologous sequence tracks.

**Figure 2.**
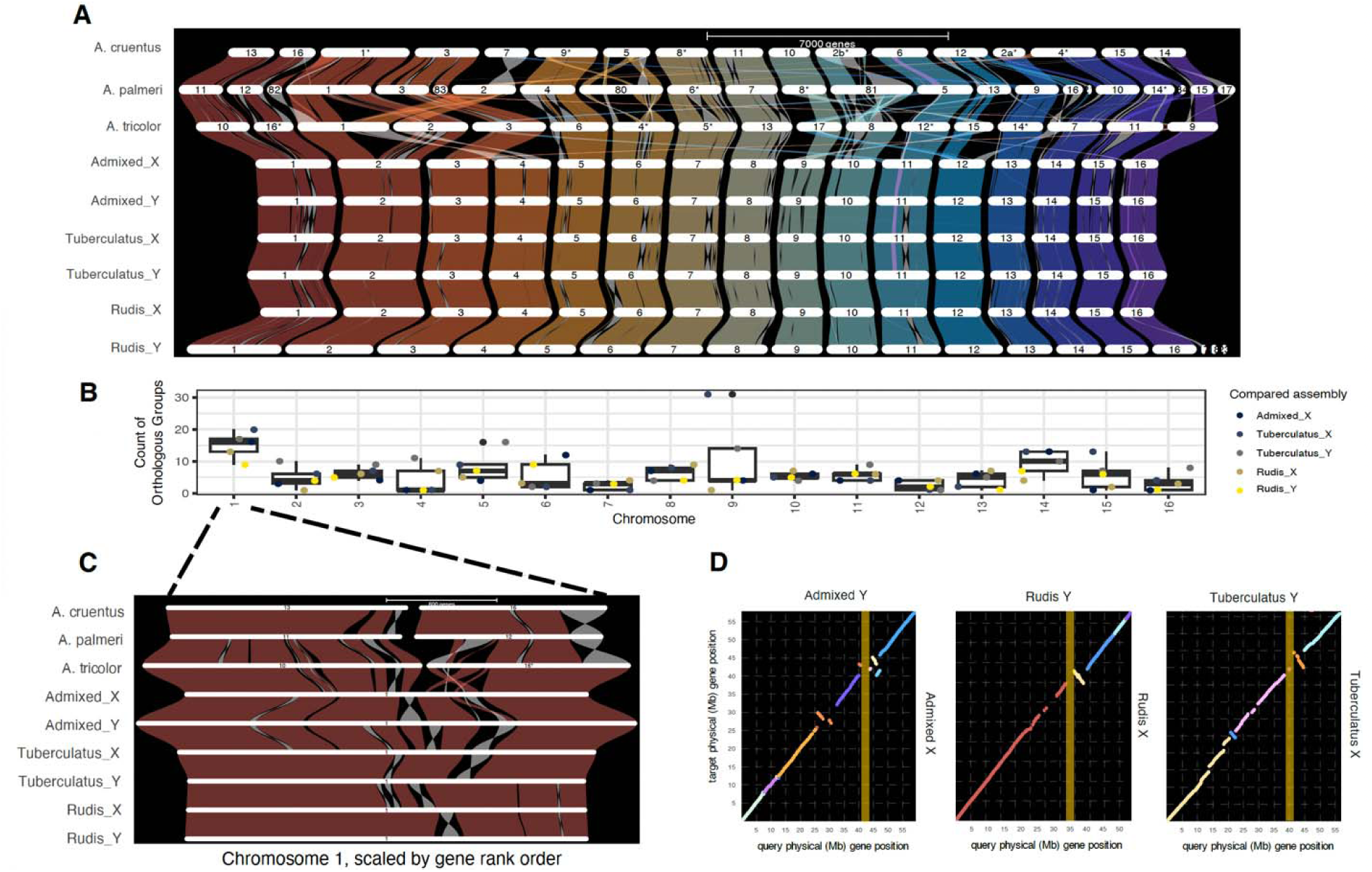
Structural variation within *A. tuberculatus* is enriched on the chromosome containing the sex-linked region. ***A***) A synteny plot from genespace highlighting the syntenic gene orthologues within *A. tuberculatus* among haplotype-phased assemblies and between *A. tuberculatus* and three congeners (*A. tricolor, A. palmeri,* and *A. cruentus*) with chromosome level assemblies. Inversions are highlighted in light grey, as are translocations which can be distinguished from inversions by their connections between distinct chromosomes. ***B***) The count of orthologous groups between the focal Y-containing (“Admixed”) assembly and each other haplotype-level, along with box plot summaries. ***C***) A zoomed-in synteny plot of Chromosome 1, which contains the SLR, illustrates the widespread occurrence of inversions ***D***) Pairwise dotplots of syntenic genes between the proto-X and proto-Y chromosome-level assemblies for each male. The SLR, highlighted in yellow in panels C and D, is shown on the focal assembly to which population genomic data was mapped.

Genotype-phenotype associations resolved a large sex-linked region, ∼3.0 Mb long, with a near continuous butte of extreme genotypic correlations with sex on Chromosome 1, haplotype 2 (but not haplotype 1 of the same genotype when a competitive mapping approach was taken; **Figure 1A**; **Sup Figure 3)**. This sex-linked region (SLR), herein referred to as the primary SLR, contained 66 annotated genes, including candidates with strong roles in sex-specific traits such as MADS8 [45] and FT/HD3 [46]. This primary sex-linked region is about 10 Mb downstream of the putative centromeric region according to RepeatOBserver (which uses a Fourier transform of DNA walks to identify putative centromere locations [47]; **Sup Figure 4**). Apart from the centromere, this region shows some of the highest patterns of 0-fold diversity and lowest gene density across the chromosome (**Sup Figure 5)**. The primary SLR shows a strong signal of highly relaxed selection, with a median πN/πs = 0.993, relative to the chromosome-wide median of 0.384.

**Figure 3.**
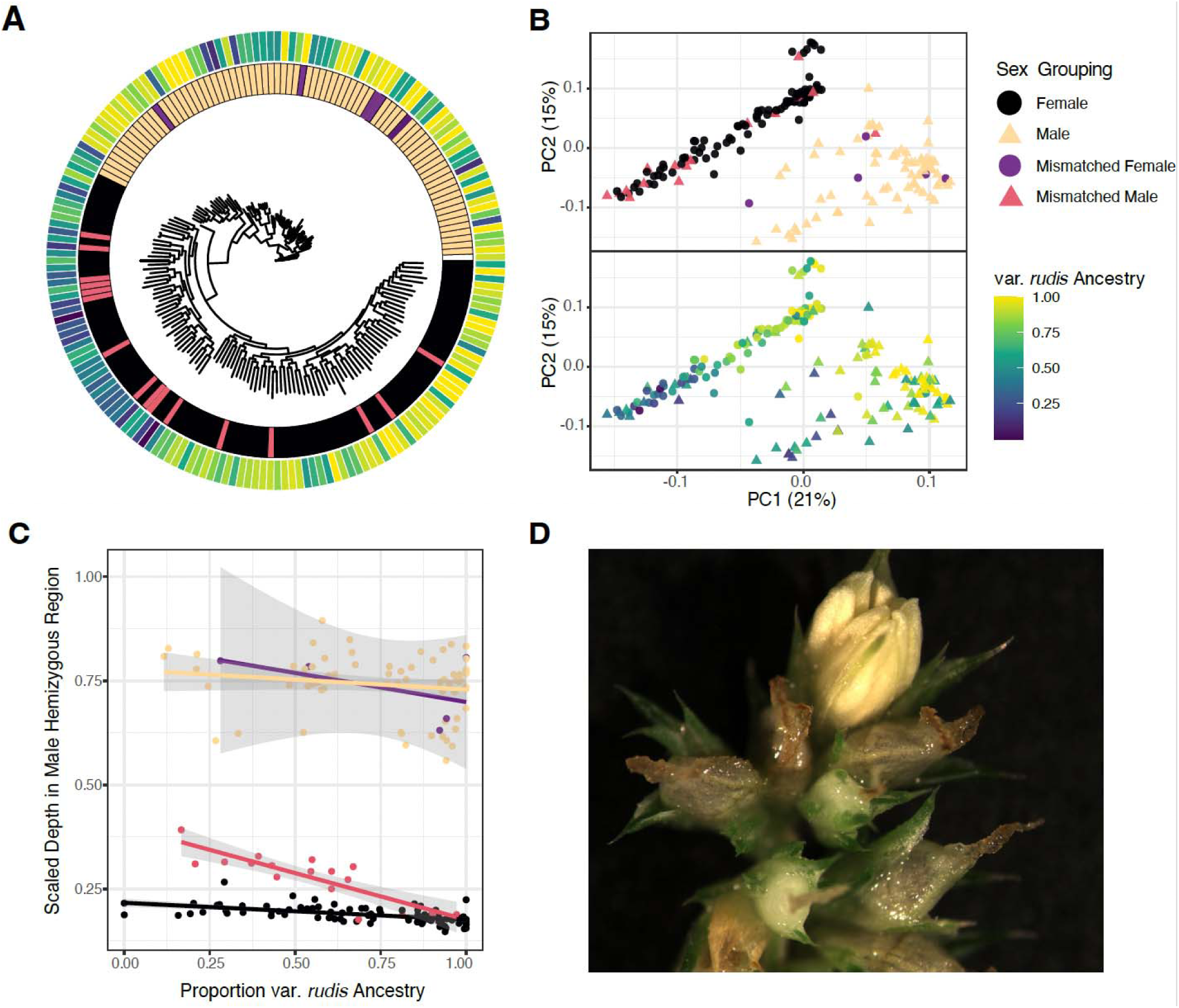
Diversity in sex at the genomic and phenotypic level in *A. tuberculatus*. **A**) A phylogeny inferred from IQtree of SNP genotypes in the sex-linked region, with tips coloured by sex grouping and var. *rudis* ancestry proportion (legend in B). **B**) PCA of SNP genotypes the sex-linked region, coloured by sex grouping and var. *rudis* ancestry composition. **C**) The median scaled depth in the SLR is predicted by the proportion of var. *rudis* ancestry, and by the interaction of sex grouping x ancestry. **D)** A male flower (emerging anthers in light yellow) atop a predominantly female inflorescence (stigmas in brown emerging from female flowers) in a waterhemp individual.

**Figure 4.**
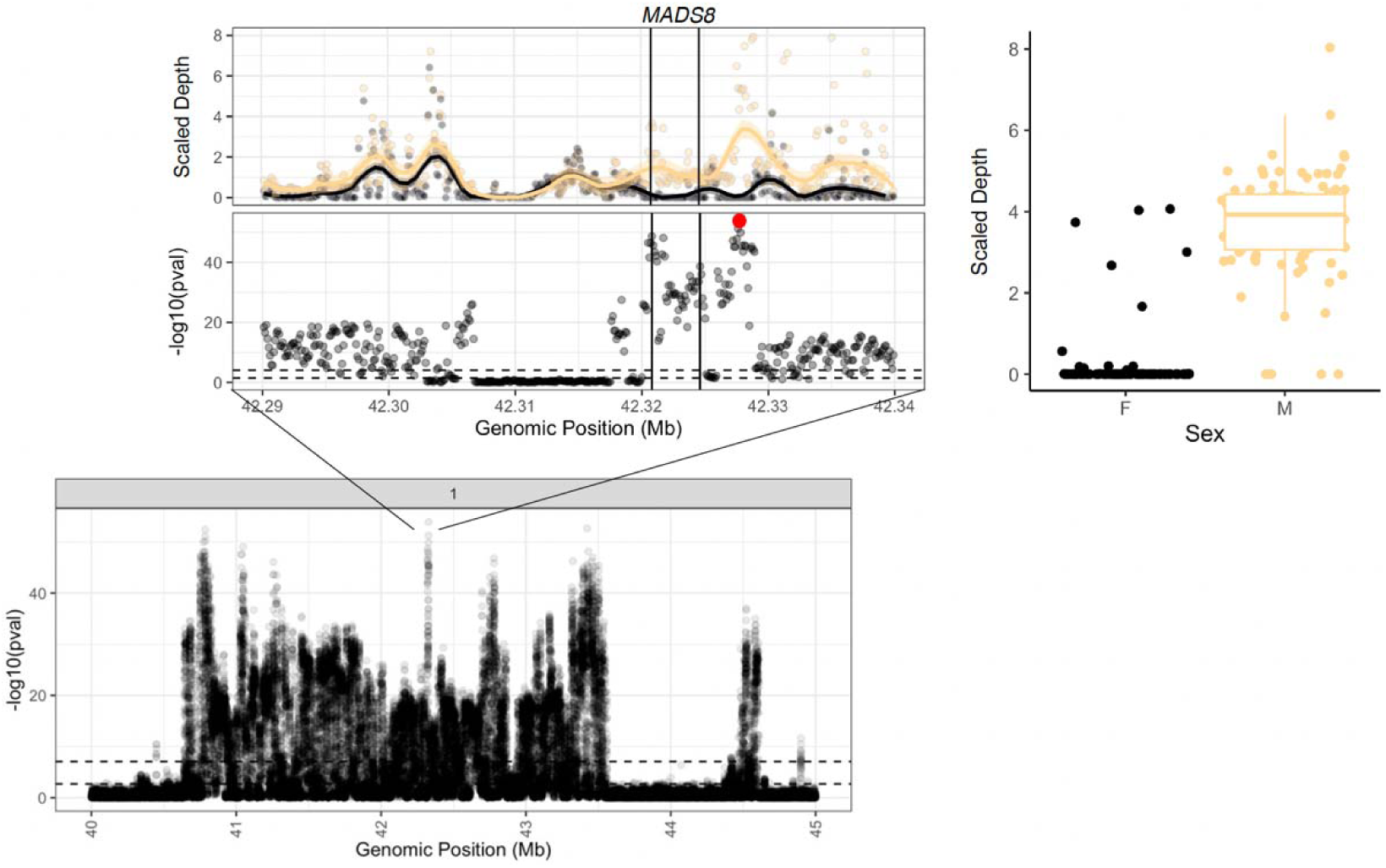
Coverage-based GWA resolves fine-scale variation in the association with sex within the sex-linked region. A) The Manhattan plot of significance of the association of sex with mean scaled depth calculated in 100bp windows across the sex-linked region. B) Zooming in on the region of the genome with strongest depth-based association with sex, [*upper*] the distribution of mean scaled depth in 100bp windows for individuals phenotyped as male (yellow) and female (black) [*upper*] and the significance profile of this depth-based variation with sex. C) Sequence content variation at the 100bp window with the strongest association with sex, highlighted in red in B).

To identify whether the primary sex-linked region was male or female specific, we calculated read depth across the region in each sample. By scaling each individual’s median read-mapped depth in genomic windows within the sex-linked region by the median read-mapped depth on Chromosome 1 outside of the sex-linked region, we identify a clear signal of male-specific hemizygosity in the region. While median male and female scaled-depth show complete overlap for most of Chromosome 1, there is clear divergence among sexes in sequence presence/absence between 40.5-43.5 [and a smaller secondary peak around 44.5] Mb. Most males show a median scaled-depth around 0.75 on average across this region (light yellow in **Figure 1B**), while most females (black in **Figure 1B)** show a median scaled-depth closer to 0.25. This is distinct from the expectation of a fully hemizygous Y-specific region, for which the median scaled depth should hover around 0.5 and 0 for males and females respectively. This pattern may reflect the presence of duplicated regions within the putative Y-specific region, as revealed through investigations of non-orthologous synteny in the proto-Y vs proto-X chromosome (**Sup Figure 6),** which may enable mis-mapping of reads from elsewhere in the genome into this region. Regardless, coverage-based, GWA, F_ST_ and heterozygosity analyses (**Sup Figure 7**) all identify the same region on chromosome 1 as the likely SDR. A synteny comparison of haplotype 1 (the proto-X) and haplotype 2 (the proto-Y) of Chromosome 1 assemblies supports these population-level inferences, revealing that the Y sex-linked region lacks syntenic orthologous genes found on the proto-X (**Figure 1C**).

The sex GWA also showed several significant associations (surpassing a Bonferroni threshold of α=0.05) outside of the primary SLR on and off of Chromosome 1 (**Figure 1, Sup Figure 1**). Notably, a locus ∼8 Mb upstream of the primary SLR shows the strongest sex association genome-wide (**Sup Figure 8**), falling largely within the gene *Agamous-like MADS-box* (AP1). Four SNPs in this peak are non-synonymous mutations, encoding an amino acid substitution in the protein. Analysis of Hi-C contact data for this sex-associated locus, along with a second locus located ∼1 Mb downstream of the primary SLR, supports their assignment to the proto-Y assembly (**Sup Figure 9, Sup Figure 10).** Their association remains even when multi-mapping reads are excluded (**Sup Figure 2**). We tested the redundancy/information value of these sex-linked loci outside of the primary SLR and off of Chromosome 1 using a lasso regression approach after the exclusion of multi-mapping reads (**Sup Figure 11**). For the 658 sex associated loci across the genome that pass a Bonferroni correction of q < 0.05 and have no missing data, allowing for the optimal shrinkage of predictors results in 42 loci with non-zero effects on sex, for which 27 are found on Chromosome 1, and 16 are within the primary SLR. This suggests that reads mapping to these genome-wide regions explain independent variation for sex. However, the r^2^ of the lasso model still remains at 88%, suggesting remaining unexplained variation in sex phenotypes.

To determine whether sex-associated loci outside the primary SLR are artifacts of mapping biases due to alignment against a single, linear reference genome, we compared male-to-female coverage ratios (indicative of hemizygosity) of sex-associated SNPs genome-wide to those within the primary SLR. Among the 864 significant sex-linked SNPs found off of Chromosome 1, male:female coverage was elevated (mean ratio = 1.20, 95% empirical range: [1.01,1.52]), their distribution largely overlapping with that of sex-linked SNPs on primary SLR (mean = 1.49, 95% range: [0.99,2.59]) (**Sup Figure 12**). The 20% enrichment in male-to-female coverage of off-Chromosome hits suggests that this proportion of males may possess sex-linked sequences on their Y haplotype that are absent from our focal Y reference, leading to mis-mapping elsewhere (as in [48]]). This finding indicates that *A. tuberculatus* Y haplotypes may be quite diverse, challenging single-reference approaches, and the prediction that Y chromosomes are degenerate and low in variation [49,50].

### Structural variation across male waterhemp genomes and the sex-linked region

To investigate the scope for Y haplotype variation further, we generated between 34.7 and 45.7 Gbp of PacBio HiFi data and between 19.6 and 21.9 Gbp of chromatin conformation capture (Hi-C) data for two additional male *A. tuberculatus* plants from across its range. We used these data to generate haplotype-resolved genome assemblies for each plant. The genome assemblies each contain 16 large scaffolds representing the 16 chromosomes of *A. tuberculatu*s along with many small scaffolds that could not be incorporated into the main assembly (**Sup Table S1**). The six haplotype-phased, chromosome-level assemblies we generated provide the opportunity to further resolve the extent of structural variation within the *Amaranthus* genus (**Sup Table 1)**. Consistent with recent reports [42], our analyses show that the largest chromosome in *A. tuberculatus* (1) was formed through a chromosomal fusion (i.e. Robertsonian translocation) since the split of *A. tuberculatus* from *A. tricolor, palmeri,* and *cruentus* (**Figure 2A)**.

Our data also illustrates widespread intraspecific structural variation in *A. tuberculatus*. Each genome shows signals of an ancient whole genome duplication as suggested in [51], with ohnologues present on distinct chromosomes (e.g. **Sup Figure 13, 14)**. When comparing the 6 *A. tuberculatus* haplotype assemblies to each other, we find numerous inversions, duplications, and translocations (**Figure 2A**). To quantify the extent of structural rearrangement, we calculated the number of orthologous groups (contiguous tracks of syntenic sequence) within each chromosome between each pair of haplotype assemblies. We found on average ∼5-8 orthologous groups per chromosome depending on the paired comparison (**Figure 2B**). The SDR containing Chromosome 1 shows a higher number of orthologous groups than nearly all other chromosomes in a univariate regression (chromosome effect: F = 2.9352, P = 0.00157; pairwise t-tests: P < 0.05 for all comparisons except Chromosome_9). This effect is especially noticeable in the high number of inversions segregating across these haplotypes (n = 11, **Figure 2C, 2D**), compared to the number of orthologous groups observed in each pairwise comparison (lower and upper quartiles=13, 20). When controlling for chromosome length in a multivariate regression, Chromosome 1 no longer appears as an outlier for the number of orthologous groups. This suggests that the structural complexity observed in Chromosome 1 may be a function of its size rather than its role in sex determination.

Given the number of structural rearrangements present on the chromosome containing the primary SLR, and the well-documented role of structural variation in limiting recombination along SLRs [52–54], we investigated whether these rearrangements could be sex-linked. This analysis is particularly relevant as Raiyemo et al. [42] recently reported two large inversions within chromosome 1 in their study of sex determination in waterhemp, though the extent of their sex-association remained unclear. To understand the role of inversions in sex determination, we asked whether there was sequence consistently inverted between X- and Y-containing haplotypes of Chromosome 1 using genespace [55]. To do so, we first assigned haplotypes as X or Y based on the expected excess of sequence content in the SLR in Y compared to X haplotypes. While a number of private and low frequency inversions are present, segregating between just X-containing assemblies (**Sup Figure 15**) and between just Y containing assemblies (**Sup Figure 16; Sup Table 2**), we see no inversion consistently inverted between the X and Y. One notable inversion is directly upstream and neighboring the sex-linked region, but is polymorphic in only ⅔ X-Y comparisons and lacks any notable signal of sex-linkage based on population genomic inference (i.e. GWA in **Figure 1**; F_ST_ and Heterozygosity in **Sup Figure 7**).

We revisited our population genomic data to further test whether any of these inversions exhibited cryptic population structure by sex or other variables, using PCA analyses of SNP genotypes within the boundaries of each inversion. Sex does not predict PC1, PC2, or PC3 for any of the inversions on Chromosome 1 (**Sup Figure 17**). Other variables do: ancestry (proportion of *var. rudis* ancestry based on K=2 grouping of a structure plot) loads onto PC2 of Inversion 3 and PC1 of Inversion 12; longitude onto PC1 of Inversion 11, PC1 and PC3 of Inversion 12; habitat type (Natural or Agricultural) onto PC3 of Inversion 8; and latitude onto PC2 of Inversion 8 (**Sup Table 2**). These results suggest that inversions are not facilitating population-level recombination suppression in or around the sex-determining region, but that inversion frequency may relate to differences in evolutionary history across ancestral lineages, geographic clines, and fine scale differences in habitat type.

In contrast to the broader patterns of structural variation observed across the genome, the sex-linked regions on Chromosome 1 show distinct differences in the conservation of genic content. When we compared the proportion of core genes (in this case, present in all three Y-containing assemblies) across all sex-linked loci on Chromosome 1 to genome-wide loci, we found a strong significant difference: 62% of genes were core in genome-wide comparisons (15,066/24,173), but only 8% of genes (6/71) were core within Chromosome 1 SLRs (χ² = 87.36, df = 1, p-value < 10^−15^). Notably, AP1 was among these core sex-linked genes. Gene-presence absence variation is also visually apparent in the pairwise comparisons of Y haplotypes, for which orthologous synteny is variable within the sex-linked region (**Sup Figure 18).** Some of this variation is likely due to gene gain on the Y or loss on the X, as 3/71 sex-linked genes on Chromosome 1 are present on the focal Y but absent on all other X assemblies. Much of this variation may relate to the repeat structure of genes within the SLR, as numerous paralogues of sex-linked genes are found elsewhere in the assembly (**Sup Figure 5**). However, since repeat-rich, low-gene-density regions are among the most challenging to assemble [56], technical limitations in assembling the Y region may also contribute to the presence/absence variation we observe. This is especially a limitation for the tuberculatus haplotype assemblies, as we found that running HiFiasm without Hi-C data initially, and only later using it for scaffolding, was necessary to avoid highly asymmetric haplotype sizes.

### Genotype-phenotype mismatch and population structure in the sex-linked region

While in many systems, a single locus is sufficient to explain nearly all phenotypic variation in sex, this seems not to be the case in *A. tuberculatus.* We see a strong notable exception to the excess of male: female coverage in the SLR for ∼10% of individuals in this dataset, which show a mismatch between sex and their average scaled depth in the SLR (**Figure 1B**; **Figure 3**). This genotype-phenotype mismatch is apparent not only in these depth profiles (**Figure 1B**, **Figure 3C)** but also with PCA and phylogenetic reconstruction of unphased genotypes in this region, for 5/94 (5.3%) and 17/89 (19.1%) of individuals phenotyped as females and males respectively (**Figure 3A, B)**. This proportional difference in genotype-phenotype mismatch represents a significant sex-bias in the lability of sex-determination (χ2=8.21, df=1, p-value < 0.0042). We also see such a signal of mismatch in sex-associated locus (AP1) upstream of the primarily SLR, with 6 females and 2 males showing mismatch, notably, with the opposite sex-bias. To test whether the degree of genotype-phenotype mismatch (the difference between an individual’s mean scaled depth in the SLR and the mean across respective sex-matched individuals) showed differences among populations, we visualized its distribution on a map and looked to identify predictors of variation in mismatch in multiple regression framework. The degree of mismatch shows a widespread geographic distribution, with mismatched individuals present in most populations (**Sup Figure 19**). While latitude, longitude, habitat type, and ancestry do not predict the degree of mismatch, U.S. state is a significant predictor (F_4,178_ = 2.4607, p = 0.047).

In response to the observation of genotype-phenotype mismatch, one might expect that individuals with a significant portion of their sex-linked region (SLR) showing genotypes associated with the oppositely phenotyped sex would also show intermediate sex expression. Alternatively, one might consider the possibility of whether sex was mis-phenotyped. To consider these two possibilities, we first independently validated our findings of genotype-phenotype mismatch with a dataset of sexed herbarium sequenced samples for which phenotypes can be reexamined, finding that again, nearly 10% of samples show a genotype-phenotype mismatch (with the same sex-bias in lability; **Sup Figure 20)**. Neither the herbarium nor the common garden population genomic datasets provided evidence of intermediate sex expression. We also tested whether genotype-phenotype mismatched individuals were more likely to have life-history trait measures (biomass and flowering time) more typical of the opposite sex, as might be expected if these individuals represent sex intermediates. Leveraging common garden phenotypes from these same samples previously ascertained [40], we tested this in a mixed regression framework (see methods), but found no such pattern (**Sup Figure 21**).

However, careful observation of reproductive structures in other contexts revealed such intermediacy, which we revisit and report here for the first time. As pictured in Figure 3D, we discovered an individual from a separate accession and grow-out that was composed entirely of female flowers, except for one male flower atop an inflorescence. In separate instances, we have observed other instances of leaky sex expression in *A. tuberculatus*, including entire branches with flowers of the opposite sex compared to the rest of the plant, and even a predominantly female plant with several perfect (hermaphroditic) flowers (though no photographs were taken at these times). This variation in sex expression supports the potential involvement of sex modifiers (regulatory, epigenetic, or environmental) that may act in the system.

Multivariate regression analyses of two distinct sex-linked metrics reveal strong population structure in the SLR (**Figure 3**). Mean scaled-depth in the SLR is structured not just by sex grouping (i.e. referring to phenotype by genotype status for each individual [M-M=male, M-F=mismatched male, F-F=female, F-M=mismatched female]; F_3,173_ = 1867, p-value < 2×10^−16^), but also by var. *rudis* ancestry (F_1,173_=7.38, p-value = 0.0073), the interaction between ancestry and sex grouping (F_3,173_=4.38, p-value = 0.0054), and marginally by habitat type (agricultural or natural; F_1,173_ = 3.3849, p-value = 0.067) (**Figure 3C)**. Genotypic structure in this region depicts an even stronger role of ancestry in shaping sex-linked haplotypes (ancestry: F = 59.6, p-value = 8 × 10^−13^; the interaction between ancestry and sex grouping: F_3,173_=3.5, p-value = 0.016), in addition to a main effect of sex grouping (F_3,176_ = 33, p-value < 2×10^−16^), latitude (F_1,173_ = 15.75, p-value = 0.0001), and habitat type (agricultural or natural; F_1,173_ = 4.17, p-value = 0.043). Clearly, the sex-linked region in *A. tuberculatus* retains plenty of diversity that reflects both deep and recent evolutionary history.

Despite the average genotype in the SLR and upstream AP1 gene being commonly mismatched to phenotypic sex, we sought out fine-scale locus-specific presence/absence variation that remained unique to one sex—a strong candidate for a SDR. To do so, we performed a GWA for sex using the scaled sequence depth in 100 bp windows as predictors, which further resolved heterogeneity in sex-linkage even within the primary SLR. While presence/absence variation at the edges of the SLR tend to show the strongest fidelity between sexes, several windows within a 10 kb tract around 42.32 to 42.33 Mb show very strong associations with sex despite being surrounded by a valley of significance (**Figure 4**; r^2^, p-value for top presence/absence locus =0.73, 1.2 × 10^−54^). Just downstream of this locus lies the MADS-box transcription factor 8, SEP1, necessary for petal, stamen and carpel development in *Arabidopsis* [57]. The median scaled depth at this upstream 100 bp window is 4 in males and 0 in females, although males show considerable variation (between 0 - 8). The genotype to sex association at this locus is still imperfect, with a fewer number of mismatched individuals, only 4 males and 7 females resembling the profile of the opposite sex (∼5% percentage mismatch), resembling the genotypic mismatch profile of AP1.

While multiple loci or modifiers may interact to determine sex, this architecture could also promote the maintenance of diversity in the sex-linked regions by allowing for gene exchange. To test this, we first performed LD based inference of the effective recombination rate (ρ=*4N_e_r;* using LDhat [58]) along Chromosome 1 (**Sup Figure 22, 23**). ρ shows a clear reduction at the border of the sex-linked region relative to the rest of the chromosome (Ratio of ρ_Y_:ρ_Chr1_ = 0.5855171). The smoothed moving average does not go to zero, and further shows a peak in ρ in the center of the SLR. It is important to note, however, that despite constraining our inference to males, that the excess mapping of reads to the Y (as observed above) will lead to an overestimation of ρ due to recombination occurring in the X and autosomal regions. The relatively low between-sex F_ST_ in the SLR might also support on-going gene exchange: while elevated compared to the genome-wide background, the median F_ST_ of 0.04 across this 3Mb region could be maintained with rare but on-going gene exchange events (based on the simplifying assumptions of F_ST_ ∼ 1/1+4N_e_*r*, where N_e_ ∼ 100,000). Along with the observation of sex-linked variation not consistently aligning with its respective sex at fine scales–even within genotype-phenotype matched individuals (**Fig 4**)—these results suggest gene exchange may contribute to observed variation in the SLR, though parallel sequence evolution through local contraction/expansion events cannot be ruled out.

## Discussion

This work demonstrates a proto-XY system in *A. tuberculatus* that retains considerable complexity and diversity despite evidence of extended linkage of loci associated with sex. We find that a complex presence/absence polymorphism between the proto-X and proto-Y is involved in sex determination. However, we find no locus that fully explains sex as phenotyped. Nearly 10% of individuals phenotyped as male or female display the genotype of the alternate sex in the SLR. Together with observations of leaky sex expression in the species, our results implicate the presence of additional sex-modifying factors and the maintenance of diversity in primary SLR.

We used long-read sequencing technology in combination with Hi-C sequencing to generate haplotype-resolved genome assemblies for three unrelated male waterhemp plants. Through GWAS, we identified a ∼3 Mb region strongly associated with sex on the longest chromosome near a chromosomal fusion junction. We also found additional sex-associated loci both upstream and downstream of this region on the same chromosome. Hi-C contacts confirmed the correct phasing and orientation of contigs placed within this region (**Sup Figure 9**), however several gaps remain in the assembly of the primary SLR (**Sup Figure 10**). In addition, we identified several much smaller sex-associated loci on other chromosomal scaffolds that remained sex-associated even after stringent filtering for mapping quality and multi-mapping read status, although most are collinear with SNPs within the SLR. One explanation for these signals may be that considerable structural variation across the sex-determining region, or gaps in our assembly, have resulted in an incomprehensive reference of a canonical single sex-determining region.Reads from the sex-determining region unrepresented in our primary assembly may have therefore mapped best elsewhere in the genome. Given the complexity of structural and sequence-content variation associated with sex in waterhemp, we believe that pangenomic approaches will be crucial for fully resolving sex-linked genomic diversity.

The evolution of dioecy from monoecy has classically contrasted pathways that rely on large or small effect mutations [1]. Recent models on the gradual evolution of separate sexes through selection on sex allocation however outline how polygenic variation for sex allocation can be concentrated to a single locus through ecological selection for sex specialization [14]. Under such a model, XY systems are more likely to evolve when fitness exponentially relates to female allocation, which could result from the coupling of survival and seed production in highly fecund females. Such a scenario is plausible in *Amaranthus tuberculatus*, as females are estimated to produce between 35,000-1,200,000 seeds per plant [59], contributing to the challenge of managing waterhemp invasions in agricultural fields. While we identify large-effect loci capable of explaining either 82 or 76% of the phenotypic variation in sex (depending on SNP or presence/absence variation), no locus shows a perfect correlation with sex. Thus, despite *A. tuberculatus* exhibiting a significant concentration of the genetic architecture of sex within a ∼3 Mb region on the largest assembled chromosome, this work implies the persistence of segregating variation. The integration of expression data with GWA studies (i.e. eQTL studies [60]) holds significant promise for uncovering potential modifiers of sex expression in the species.

The evolutionary and genetic mechanisms underlying sexual intermediacy in dioecious species are thought to vary from unstable (developmental plasticity–epigenetic), to transitory (paradioecy–polygenic) or selectively reinforced (leakiness)—although genetic mechanisms are unknown in most systems (discussed in [22,26]). In *Silene*, which is known to have evolved through the two-step pathway, variation in the expression of sex is influenced by epigenetic regulation [61]. Extensive phenotypic observation of herbarium samples in the *Siparuna* genus revealed “inconsistent” sex expression in males and females, despite the lack of such observations in contemporary samples, providing support for the paradioecy model [62–64]. In *Mercurialus*, recent experimental and theoretical work has demonstrated that disruptive selection on sex allocation drove the evolution of dioecy [18], with subsequent selection on mate-limitation maintaining variation in its expression [24,65]. For the primary SLR, we find lability in sex-determination to be greater in males as compared to females in multiple datasets (also observed, albeit at a lower rate by [66]), which would be consistent with stronger mate-limitation in males acting to maintain variation for sex-expression. While we document a single case of inconstant sex expression, the scale of our phenotyping—nearly four thousand individuals assessed for sex and other traits in a common garden [40]—suggests that subtle variation in sex expression may have gone undetected. Notably, the congener *A. palmeri* exhibits transient hermaphroditism, where staminate (male) flowers initially develop both male and female reproductive organs [67], suggesting that the developmental pathways enabling hermaphroditism may still be segregating in recently evolved dioecious *Amaranthus* species. Future work in *A. tuberculatus* will look to resolve the frequency of inconstant sex within and across populations, and integrate regulatory and epigenetic investigations (as in [68]) to decipher the genetic mechanisms maintaining this variation.

Multiple mechanisms of sex determination might be selectively favored to prevent the degeneration of sex-linked regions [69]. The primary sex-linked locus shows evidence of substantial presence/absence heterogeneity both within and among sexes, with sequencing depth in 1 kb windows varying between 0 - 39.3X for males and 0 - 13.5X for females. When this fine-scale variability is associated with sex, it displays a characteristic “suspension bridge” pattern typical of recombination suppressed regions, with divergence concentrated near the breakpoints and troughing at the center of the region where selection counters recombination [53,70,71]. The peak of *4NeR* in the center of the sex-linked region, along with a 10% rate of phenotype-genotype mismatch, could be interpreted as evidence for on-going gene exchange (i.e. “flux” through interlocus gene conversion, double- or successive recombination events) countering sex-specific selection [72,73], similar to recent observations for inversions encoding wing coloration in *Heliconius* [72]. Gene conversion among sex chromosomes has been recognized as a mechanism maintaining variation in sex chromosomes in several species, such as *Drosophila* and humans [73,74]. Alternatively, these patterns might result from independent sequence evolution through local contraction/expansion events in repetitive regions of both X and Y chromosomes, creating parallel changes that mimic the signatures of genetic exchange. If genotype-phenotype mismatches were primarily driven by gene exchange events, we might expect a geographic clustering of mismatched individuals, reflecting the historical spread of recombinant haplotypes. We find mixed evidence for a geographic pattern, with mismatched individuals distributed across nearly all populations but enrichment in particular U.S states (**Sup Figure 19**). Distinguishing between these alternatives will require fine-scale recombination rate maps across the sex-linked region and population-level haplotype reconstruction of Y chromosomes across the species range.

There is clear evidence for the maintenance of variation in sex-linked haplotypes across the native range of *A. tuberculatus.* These results parallel the remarkable diversity among Y haplotypes that has been recently observed in guppies and fruit flies [48,75][73,74]. We find that diversity in sex-linked haplotypes is structured by ancestry, geography, and habitat. On one hand, this pattern may reflect neutral divergence, with sex chromosomes diverging on either side of the Mississippi River before the secondary contact between *Amaranthus tuberculatus* var. *rudis* and var. *tuberculatus* [59]. Alternatively, it could result from climatic selection. Our previous analyses demonstrated an interaction between sexual and natural selection on flowering time, with males exhibiting stronger divergence from females at lower latitudes [40,41]. This pattern suggests that divergent sex-linked haplotypes may play a role in local adaptation.

Our analyses reveal two key MADS-box transcription factors associated with sex determination in *A. tuberculatus*. The narrow peak of sequence presence/absence association in the middle of the SLR corresponds to MADS8 (SEP1), where the relative dearth of associations around it suggests particularly strong sex-specific effects [76–78]. Separately, the peak with the strongest genotypic association with sex, located ∼8Mb upstream of the primary SLR, falls within an Agamous-like MADS-box transcription factor, AP1. The presence of two MADS-box genes at these distinct sex-associated loci supports their convergent role in regulating sex differentiation [79]. MADS box transcription factors play an exceptional role in developmental novelties, with their duplication and neofunctionalization in plants resulting in sex-specific development, flowers, fruits, and seeds ([80–83]), and in *Populus*, potentially even the transition from male to female [84]. MADS8 in particular has been characterized as a gene indispensable for female development in Rice (*Oryza sativa*) under high temperatures [45]. Such an environmentally-dependent pathway could explain the lack of complete phenotype-to-genotype mapping we observe for MADS8. Previous work on the genetics of sex in *A. tuberculatus* identified *Flowering Locus T* (*FT*) as a candidate for a sex-determining gene, having been shown to be sex-linked and differentially expressed across male and female flowers throughout development [31,42]. While *FT* is present in our primary Y assembly, it shows presence/absence variation across our three Y assemblies, which could reflect either true biological polymorphism among Y haplotypes or technical artifacts from assembly challenges in this complex region. With genetic transformation protocols advancing in the species [85], future work will look to distinguish whether these sex-linked genes play a functional role in sex determination.

These results provide support for the concentration of the genetic architecture of sex during the transition from monoecy to dioecy. While large-effect alleles have accrued on the extended sex-linked region on Chromosome 1, the genetic architecture of sex and its expression is variable. This variability appears to be facilitated by either on-going gene exchange between the proto-X and Y or parallel sequence evolution through local contraction/expansion events. In either case, the absence of a fully sex-limited locus suggests modifiers of sex facilitating lability in its expression. From an applied perspective, this complexity poses a significant challenge for strategies such as gene drive targeting the sex-determining locus for weed control. Future work exploring the genetic basis and fitness consequences of sex allocation in closely related monoecious species (as in [18]) will be key to disentangling the ecological forces and genetic constraints that have shaped the evolution of dioecy in *Amaranthus*.

## Acknowledgements

Thank you to Tyler Kent, Sally Otto, Carl Veller, and the Rieseberg and Mank Labs for feedback on the paper. Thanks to Hayley McKay for generating the gene expression data used for gene annotation. Thanks to Federico Trucco for observing the leaky sexed waterhemp individual and permission to use the photo here.

## Data Availability Statement

Genomic sequences generated for this project have been deposited into the Sequence Read Archive under project accession no. PRJNA1271601. Data underlying analyses and figures are available along with code to reproduce these analyses at <u>10.5281/zenodo.15594570</u> and https://github.com/jkreinz/SexChromPlosBio/

## Methods

### Data Production

Based on previous analyses of population structure across populations of *Amaranthus* spanning Ontario and the Midwestern United States [40,86], we selected three populations that spanned predominantly var. *Rudis* (Nune), admixed (Walpole), and predominantly var. t*uberculatus* (Nat) ancestry. Further, we selected an individual from a maternal line in an admixed population where its sibling was previously genotyped as segregating for high EPSPS copy number (∼28 copies [86]). For each population, seed from 2 maternal lines were sown into 5 pots each in growth chambers in the Biodiversity Research Centre with day and night temperatures set to (15 °C/12hr, 25 °C/8hr). After germination, pots were thinned to a single plant, and relocated to the greenhouse in early May where they would grow under lengthening days to maximize tissue production and delay the time to flowering. From each line, we collected expanding leaf tissue from one male numerous times over the summer to maximize material for high molecular weight DNA extractions and Hi-C.

Hi-C libraries were prepared using a protocol based on the protocol of Hirabayashi et al. [87]. In brief, young leaves were collected and flash-frozen in liquid nitrogen. Approximately 0.75-1 g of tissue were used as starting material. Leaves were pulverized using a mortar and pestle, and nuclei were cross-linked in 1X PBS buffer + 1.5% formaldehyde for 15 minutes at room temperature in a rotisserie oven. Cross-linking was halted by adding glycine at a final concentration of 250 mM, and incubating the tissue for 5 minutes at room temperature, with rotation. Tissue was washed once in ice-cold 1x PBS buffer and resuspended in 10 ml of cold Nuclei Isolation Buffer (20 mM HEPES pH 8.0, 250 mM sucrose, 1 mM MgCl_2_, 5 mM KCl, 40% glycerol v/v). It was then filtered, in successions, through two layers of cheesecloth and one layer of miracloth (MilliporeSigma, Burlington, MA, USA). Nuclei were then washed in Nuclei Isolation Buffer and fractionated using a 95% Percoll (MilliporeSigma) gradient buffer. DNA in purified nuclei was then treated with DpnII restriction enzyme, biotinylated, and proximity-ligated. DNA was extracted from the nuclei using phenol:chloroform:isoamyl alcohol (Invitrogen, Waltham, MA, USA) and quantified using a Broad Range kit on a Qubit fluorometer (Invitrogen). Biotinylated DNA fragments were isolated from 1-4 µg of the resulting DNA, and Illumina libraries were prepared following Todesco *et al.* 2020 [88]. Preliminary experiments performed in sunflower and cannabis showed that, due to the abundance of repetitive sequences in plant genomes, large fractions of read pairs in Hi-C libraries are not informative. Both proximally-ligated fragments in a read pair need to be mapped uniquely to the genome to provide information about long-range interactions; if one of the fragments derives from a repetitive region, and can therefore not be uniquely mapped, the read pair becomes uninformative. To reduce the abundance of repetitive regions in the Hi-C libraries, and therefore increase the proportion of informative reads, we applied an enzymatic repeat depletion treatment [88]. Hi-C libraries were sequenced by Novogene Corporation Inc. (Sacramento, CA, USA).

HMW DNA for PacBio sequencing was extracted using a CTAB-based protocol adapted from Stoffel, K. *et al*., 2012 [89] in the Rieseberg Lab at UBC. Briefly, 0.8 g of flash frozen young leaves of *Amaranthus tuberculatus* were ground to thin powder using mortar and pestle in liquid nitrogen. The powder was thoroughly resuspended in Extraction Buffer (100mM Tris-HCl pH 8.0, 1.4M NaCl, 20 mM EDTA pH 8.0, 2% w/v CTAB and 1% v/v b-mercaptoethanol) and incubated 1 hour at 55°C. Eventually the tube was cooled down to RT and extracted with one volume of chloroform:isoamyl alcohol (24:1). The aqueous phase was supplemented with NaCl up to a final concentration of 2.6 M and re-extracted with chloroform:isoamyl alcohol. The new aqueous phase was transferred to two new tubes that were topped up with at least 6 volumes of Precipitation Buffer (50 mM Tris-HCl pH8.0, 10 mM EDTA pH 8.0 and 1% w/v CTAB). Tubes were centrifuged for 30 minutes at RT and the combined pellets were washed with milliQ water. Eventually the pellet was gently and fully resuspended in NaCl 1.5 M and incubated with RNAseA for two hours at 37 degrees. After the RNAse treatment, a third chloroform extraction was followed by two washes with 75% ethanol. Finally the fully dried pellet was allowed to resuspend overnight in 10 mM Tris-HCl pH 8.0 at 4°C. Pipetting was reduced to the minimum along the procedure and if done was exclusively handled with wide-bore tips to avoid damaging DNA. HiFi sequencing was performed at the Yale Genomics Centre. We decided to sequence one individual from Walpole, Ontario, Canada, to high coverage (70X), and the other two individuals to the minimum coverage recommended by HiFiASM for haplotype-phased assembly + HiC (25X) to assess assembly quality at varying coverages [90].

### Genome assembly and annotation

Hifiasm was used to produce haplotype-phased assembly, informed by Hi-C data for each sample. Default settings were used, except for one sample (which happened to have lower HiFi coverage; var. *tuberculatus*) which showed asymmetric assembly sizes between haplotypes. As recommended in the HiFiasm documentation, the setting for heterozygosity was adjusted to multiple values, however with little effect. We found that removing the Hi-C data from the initial Hifiasm assembly for this sample led to a much more symmetric haplotype size, and thus we proceeded with this approach. After the assembly of each haplotype’s contigs, contigs were scaffolded using the Hi-C data with the program YaHS [91], and the resulting scaffolds were then manually curated with juicebox (using the ARIMA pipeline https://github.com/ArimaGenomics/mapping_pipeline) based on the contact map. Briefly, scaffolds were joined together that had overrepresentation of long-distance contacts, and contigs were flipped in orientation that showed an excess of long range compared to short-range contacts within the scaffold. Finally, we performed pairwise mapping of each haplotype to each other, and to one haplotype of each other individual, using the SynMap2 tool in Coge [92] to confirm the consistency of scaffold naming among assemblies for the 16 largest scaffolds that each represents a chromosome (such that Chromosome 1 is syntenic to Chromosome 1, and Chromosome_2 to Chromosome_2, etc., across all haplotypes). We further validated the phasing of haplotypes in particular regions of interest in our focal assembly (admixed var. tuberculatus and var. rudis). We concatenated the two haplotypes into one file and determined Hi-C contacts across the concatenated haplotypes with juicer to search for regions with contacts between haplotypes but not within a single haplotype (i.e. regions that are misphased). Assemblies quality was quantified using Quast (https://github.com/ablab/quast) and BUSCO with plants as the taxonomic grouping [1].

All six haplotype assemblies were then annotated with Maker [93], using *ab-initio* prediction, along with protein sequence from a previous *Amaranthus tuberculatus* assembly [86], its congener *Amaranthus hypochondriacus* [94], and using transcriptomic data we produced from leaf tissue from 24 individuals spanning two populations from southwestern Ontario found in a Natural and Agricultural setting, with 12 males and 12 females sequenced. For these transcriptomes, RNA was extracted using RNAeasy plant kits in the Wright Lab, and libraries were prepared and sequenced at the Sickkids, Toronto sequencing facility on Illumina single-end 100bp technology.

### Comparative Genomics

To identify orthologous groupings among assemblies and visualize their synteny, we ran Genespace [55] on all 6 *A. tuberculatus* assemblies along with three other congeners with recently produced chromosome-level reference genomes: *Amaranthus cruentus* [95]*, Amaranthus tricolor* [96], and *Amaranthus palmeri* [97]. We used the gghits and riparian_plot functions to visualize pairwise and all vs all synteny. We also investigated the number of orthogroups (defined by the presence of X-1 breaks in synteny) between each haplotype assembly and across each chromosome. To identify core genes and pangenes, we queried genes for their presence/absence across assemblies both at the genome-wide level and within particular chromosomes of interest (i.e. the proto-Y). We further limited these comparisons to assemblies we expected to contain the male SLR, and used a test of equal proportions in R (function prop.test) to test for significant differences in the proportion of core genes between the SLR and genome-wide genes. We also looked to understand synteny between non-orthologous genes, using the SynMap2 function in CoGe [92]. This allowed us to compare the mapping of focal X and Y haplotypes with respect to all annotated genes, as well as calculate dN/dS between them.

### Centromere identification and Repeat Calling

Previous work generated de-novo TE libraries [41] for a previous draft *A. tuberculatus* reference genome [86]. We used repeat masker to locate these previously characterized repeats in all 6 of the new haplotype-phased chromosome level assemblies produced here. We also ran RepeatOBserver pipeline (https://github.com/celphin/RepeatOBserverV1) to identify the putative centromeres in our assembly, as inferred from the Shannon diversity index of Fourier transforms of DNA walks, with low diversity indicating highly repetitive low complexity regions as is typically found in centromeric regions [47].

### Population genomics

We leveraged 187 resequenced individuals from 17 pairs of neighboring natural-agricultural populations collected from Michigan to Kansas (from [40]) to understand the population genomics of sex-determination. These resequenced samples were grown as a part of a much larger common garden experiment, in which 6000 individuals were grown, and ∼4000 sexed and phenotyped for numerous traits [40]. Important to this study, plants were sexed at the early to mid-stages of flowering, and so were assigned a sex based on only a small proportion of flowers. Furthermore, due to the large-scale nature of this study, and the sheer number of flowers produced by an individual, we did not make an effort to look for variation in sex expression (nor did we know it existed). As a result, we could not comment on whether any individuals showed leakiness in their sex expression. In the meantime, the Tranel lab at University of Illinois Urbana-Champaign was performing a routine grow out of waterhemp to test for herbicide resistance, and noticed the individual pictured (**Figure 3D).** In addition to the pictured female individual that harbors a male flower, the Tranel lab has observed, rarely, other instances of leaky sex expression including: a predominantly female individual with an entire branch of male flowers, and an individual that contained a perfect (hermaphroditic) flower.

We re-aligned the raw reads from this resequencing dataset using BWA-mem [98] to a haplotype 2 of the highest-coverage individual (70X in total, 35X per haplotype; an individual from an admixed population in Ontario segregating for high EPSPS copy number). After calling SNPs using freebayes (options --use-best-n-alleles 2 --report-monomorphic) and filtering SNPs (using the following expressions in BCFtools [99]: QUAL >= 30, ‘AB >= 0.25 & AB <= 0.75 | AB <= 0.01’, SAF > 0 & SAR > 0, MQM >=30 & MQMR >= 30’, ((PAIRED > 0.05) & (PAIREDR > 0.05) & (PAIREDR / PAIRED < 1.75) & (PAIREDR / PAIRED > 0.25)) | ((PAIRED < 0.05) & (PAIREDR < 0.05))’) we performed a genome-wide association study of the 11,040,109 SNPs genome-wide with sex using gemma and a minor allele threshold of 5% [100]. This GWAS identified Chromosome 1 as containing the primary sex-linked region.

As loci with significant associations with sex were found to be non-contiguous across Chromosome 1 as well as found on other chromosomes, we performed a lasso regression analysis on them to determine their co-linearity in explaining sex. To do so, we extracted the genotypes of loci with p-values < 10^−15^ and analyzed them in R with the glmnet package. We performed cross-validation to identify the value of the shrinkage penalty (lambda) that minimizes the mean-square error, before refitting the data with the best lambda. We also tested whether multi-mapping status could explain these associations outside of the primary SLR. To do so, we used samtools to remove both secondary and multi mapping reads (samtools view -F 0×904) and forced freebayes to recall SNPs only at loci with significant associations with sex in the initial sex GWA. After filtering SNPs and performing the GWA for sex as above, we found no evidence that multi-mapping status could broadly explain these associations (**Sup Figure 2**). We also investigated how competitive mapping influenced our discovery of sex-linked sites across the genome. To do so, we modified the putative-Y containing haplotype assembly of the focal Walpole (admixed) individual to also contain the X-containing haplotype of Chromosome 1. We mapped reads, called SNPs, and ran a GWAS for sex, as described above. Degeneracy of SNP substitutions was inferred using the standalone python program degenotate (https://github.com/harvardinformatics/degenotate).

We then implemented a number of population genetic investigations on the sex-linked region. PCAs were performed on particular regions of interest, including each of the inversions identified in genespace and the major sex-linked region on Chromosome 1 using plink2 (--pca). Previous work inferred the ancestry of each sample with faststructure at K=2 [40], and we used these ancestry proportions that relate to the proportion of var. *rudis* or var. *tuberculatus* ancestry in a multiple regression along with sex-grouping (with female, male, mismatched female, mismatched male as factor levels), the ancestry x sex-grouping interaction, habitat (natural or agricultural), latitude, and longitude of a sample to predict PC values of each inversion. We extracted genotype calls from males and females across this chromosome (and others) to test whether sex-linked alleles showed an excess of male heterozygosity, as would be consistent with an XY system (**Sup Figure 3)**. We did so using the --geno-count option in plink for separate runs that included either only males or only females. For each sex, we then calculated the heterozygous proportion as the number of heterozygous genotypes divided by the total number of genotypes called at that locus (i.e. accounting for missing data at each locus). We used Pi_XY_[101] to calculate weir and cockerham’s F_ST_ and diversity between and within sexes, while controlling for missing data. For both F_ST_ and the difference in the proportion of heterozygous genotypes between males and females, we used R to calculate the 95% percentile of all autosomal chromosomes using quantile(probs = c(0.025,0.975) to provide context for their distributions along the sex-linked region on Chromosome 1. We also inferred the phylogenetic tree of the sex-linked region using IQtree2 [102], on sequence representing both high quality variant and invariant sites after converting SNP calls in VCF format to phylip using vcf2phylip (https://github.com/edgardomortiz/vcf2phylip).We used an iqtree algorithm that tests the best fit of 88 DNA models (options -st DNA -m TESTONLY). The best fit model was chosen according to BIC (TVM+F+I+G4). All plotting was done with ggplot2 [103] and the cowplot [104] packages in R.

We calculated a scaled-depth estimator in 100bp windows along the sex-linked region to give insight on sequence presence/absence. Specifically, we used mosdepth [105] to estimate coverage in 100bp windows for each individual directly from mapped reads. We calculated scaled-depth as the depth in a focal 100bp window along Chromosome 1 (the SLR containing chromosome) divided by the mean coverage outside of the SLR on Chromosome 1. To visualize fine-scale copy number by sample in the sex-linked region and reduce noise in these estimates, we plotted the smoothed means of copy number in 200 kb windows. Mean scaled-depth was also calculated across the entire sex-linked region.

We first used these values to identify the number and proportion of phenotype-genotype mismatch for each sex, as well as perform a 2-sample test for equality (prop.test function in R) to examine whether there was sex bias in mismatch. We then used an individual’s scaled-depth in the SLR to test whether var. *rudis* ancestry, sex grouping (defined based on male and female, and whether or not they were genotype-phenotype matched [n=4 groups]), latitude, longitude, and habitat (natural or agricultural) were significant predictors in a multiple linear regression framework. Finally, we used scaled-depth estimates in 100bp windows to perform a GWAS for sex across Chromosome 1. This was done using a custom script, for which a linear regression of sex against scaled-depth was run for each window across Chromosome 1. The -log10(p-value) of these tests were plotted, with significance assessed relative to a Bonferroni multiple test correction threshold of 0.05.

To validate the likelihood of sex mis-phenotyping, we investigated patterns of genotype-phenotype mismatch in an independent dataset, inferred local recombination rate, and the distribution of genotype-phenotype mismatch across the range. We mapped resequenced herbarium samples [106] for which the phenotype value for sex was verified and validated based on deposited images, to our focal reference genome. Mapping was performed after merging and de-duplicated reads with DeDup [107], and rescaling base-quality scores to account for DNA damage [108]. As for our contemporary samples, we then inferred copy number based on scaled depth in the sex-linked region. On contemporary samples, we inferred the effective population recombination rate (rho) with LDhat [58], using the LDhat workflow (https://github.com/QuentinRougemont/LDhat_workflow). To do so, we used a precomputed look up table (theta = 0.01) and a subsampled VCF with 100 genotypes (50 males). Rho (4N_e_*r*) was computed between each SNP. Each point estimate of rho was plotted in the SLR along Chromosome 1, to which a loess curve was also fit, while the mean rho, and the cumulative mean rho, was calculated in 100kb windows and visualized across the genome. Since we also had phenotype data for these samples, we were able to perform a multiple linear regression analysis for two key traits (biomass and flowering time), to ask whether genotype-phenotype match status predicted trait values, while controlling for experimental treatment, habitat, geography, and sex of samples. Finally, we calculated the difference in an individual’s copy number relative to the respective sex-matched mean value. We tested whether there was a strong geographic signal in the distribution in the degree of this mismatch in a multiple regression framework, with latitude, longitude, habitat, and state as predictors. We visualized this on a map using ggmap [109] and ggplot2 [103] in R.

## Supplementary Information

### Supplementary Figures

**Sup Figure 1.**
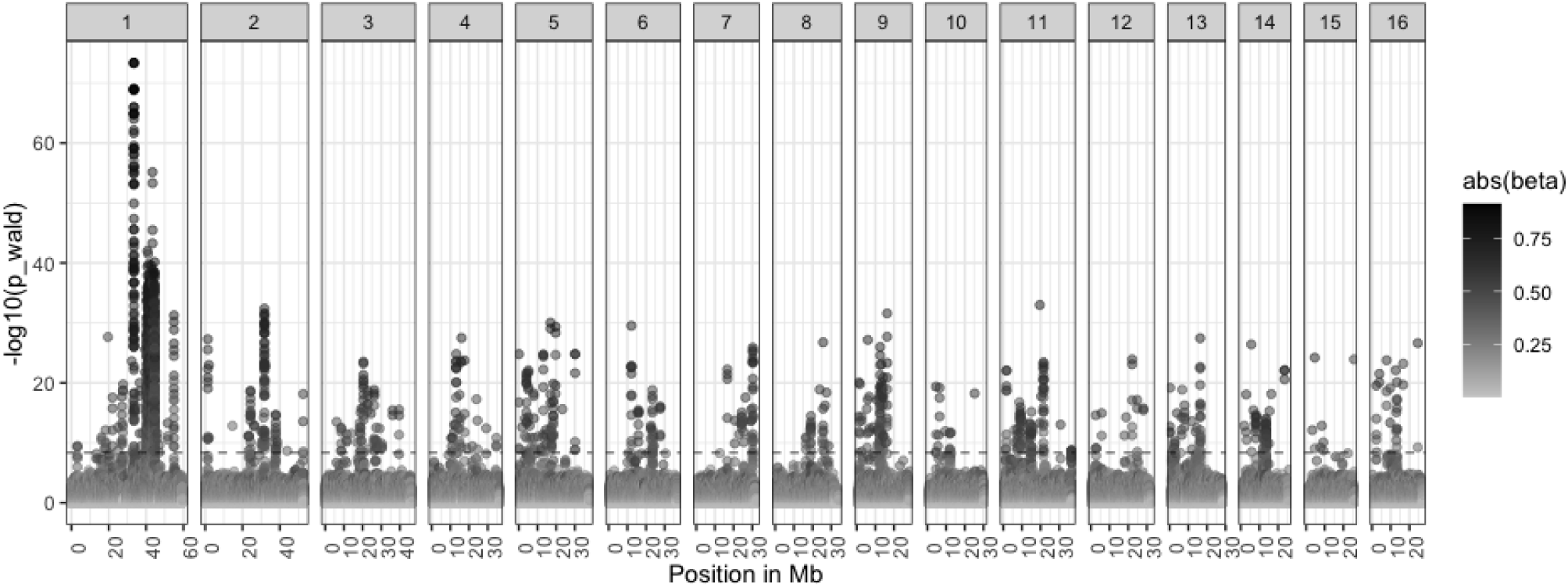
A Manhattan plot of the strength of the -log10(p-value) of the association of SNPs across the genome with sex phenotypes. Analysis was done on SNPs called from reads mapped only to haplotype_2.

**Sup Figure 2.**
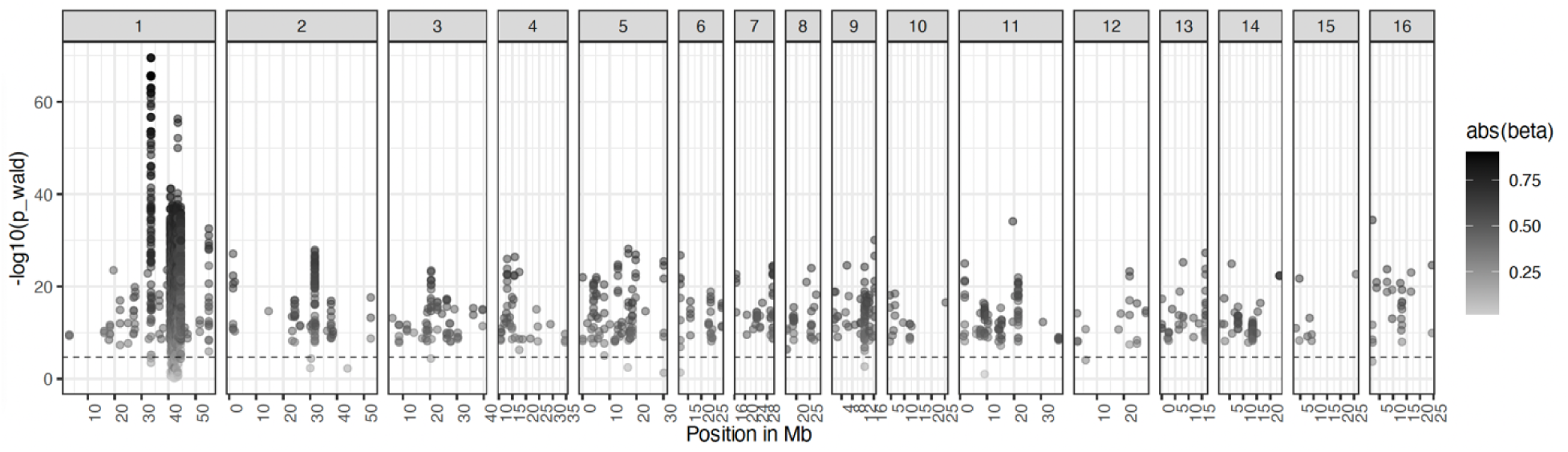
After retaining only primary alignments (removing multi-mapping and secondary mapping reads), recalling and refiltering previously ascertained sex-associated SNPs, loci around the primary SLR and off of Chromosome 1 remain significantly associated with sex.

**Sup Figure 3.**
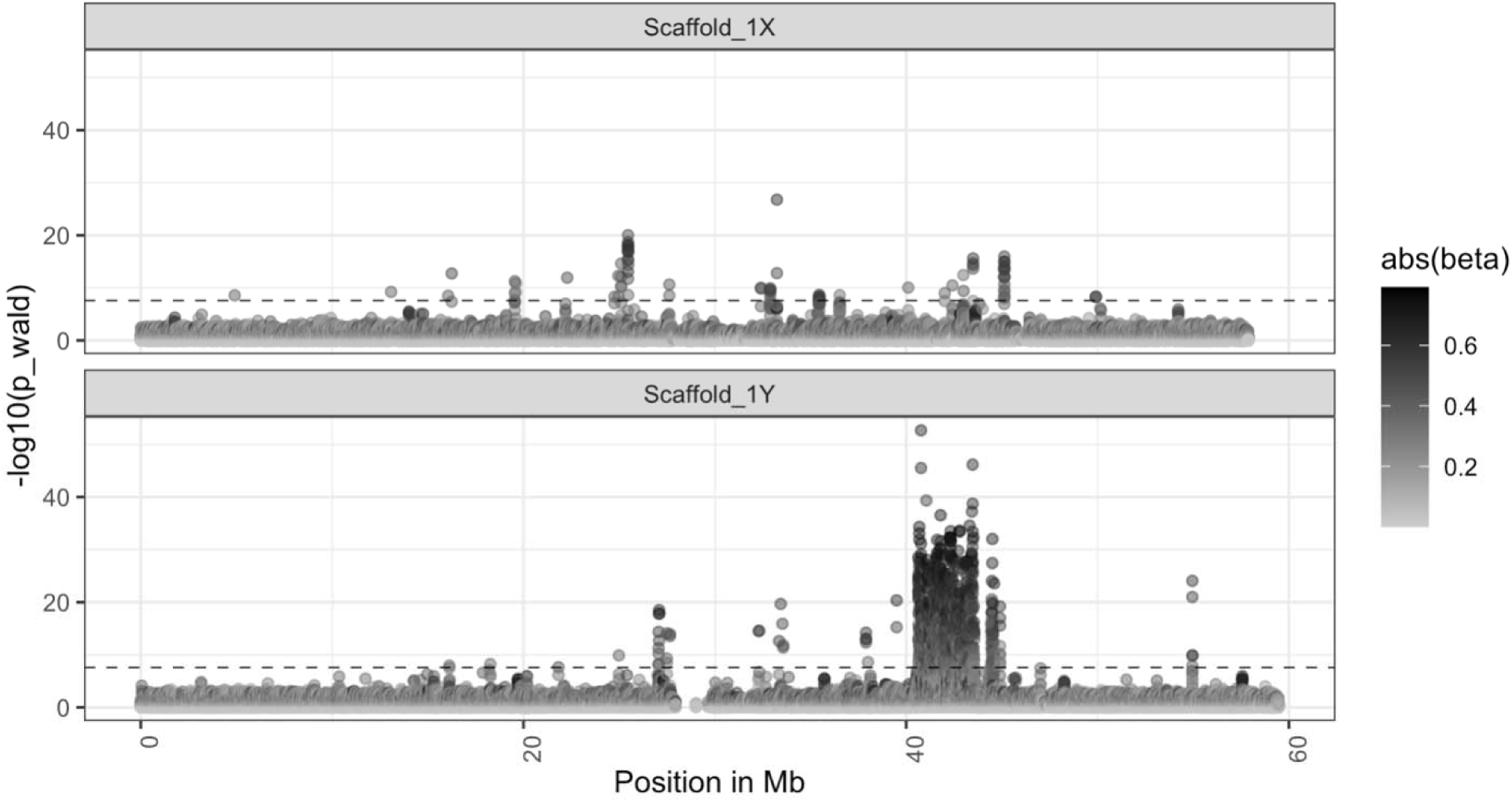
A Manhattan plot of the strength of the -log10(p-value) of the association of SNPs along haplotype_1 (the proto-X) and haplotype_2 (the proto-Y) of Chromosome 1, when SNPs are called from reads competitively mapped to haplotype 1 and 2.

**Sup Figure 4.**
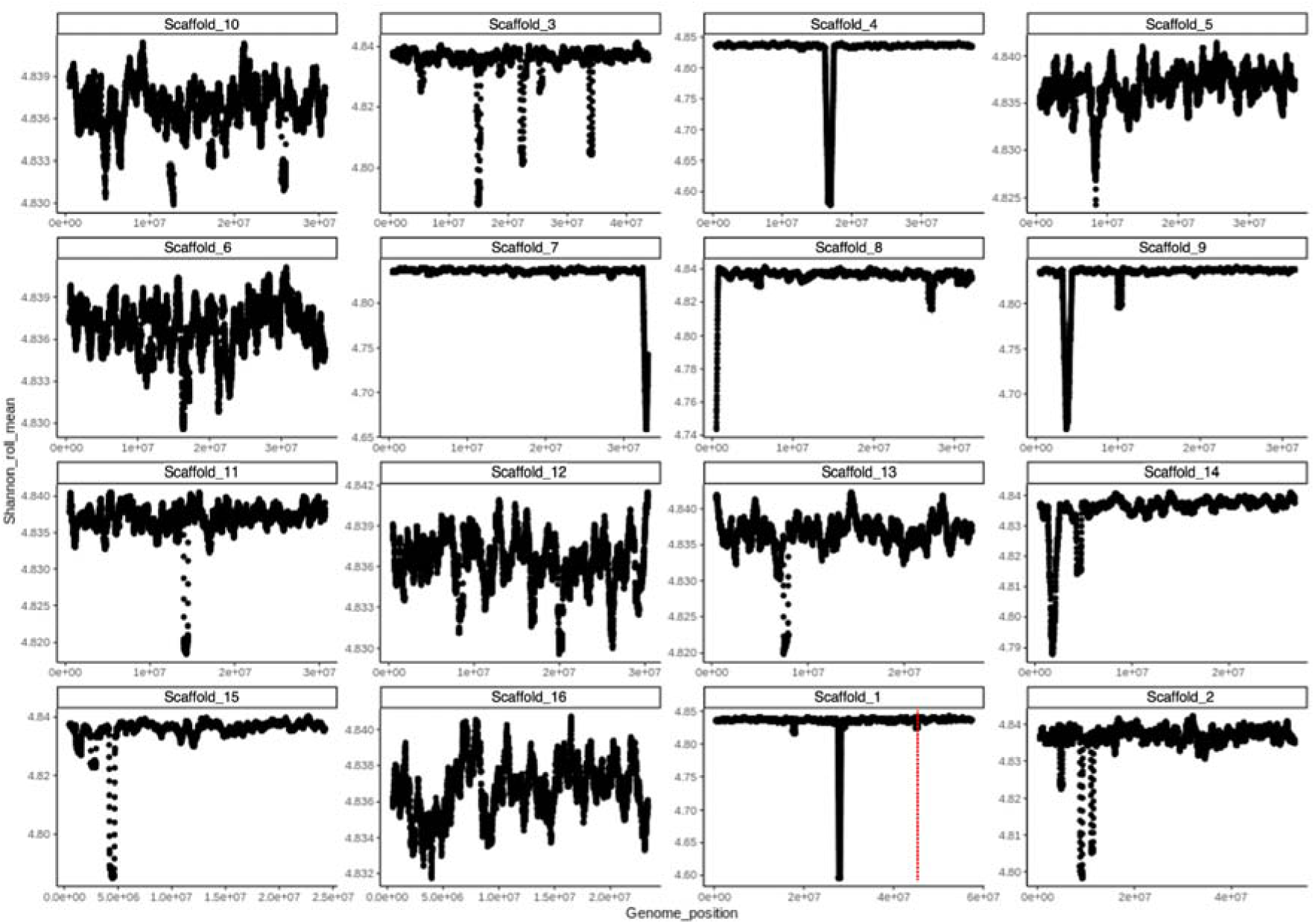
RepeatOBserver inferred shannon-diversity index of sequence complexity across each chromosome for our focal reference, where minimum values are predictive of centromere location. The sex-linked region region is present on Chromosome 1 around ∼41Mb (vertical dashed red line), and shows a tertiary minimization of Shannon diversity.

**Sup Figure 5.**
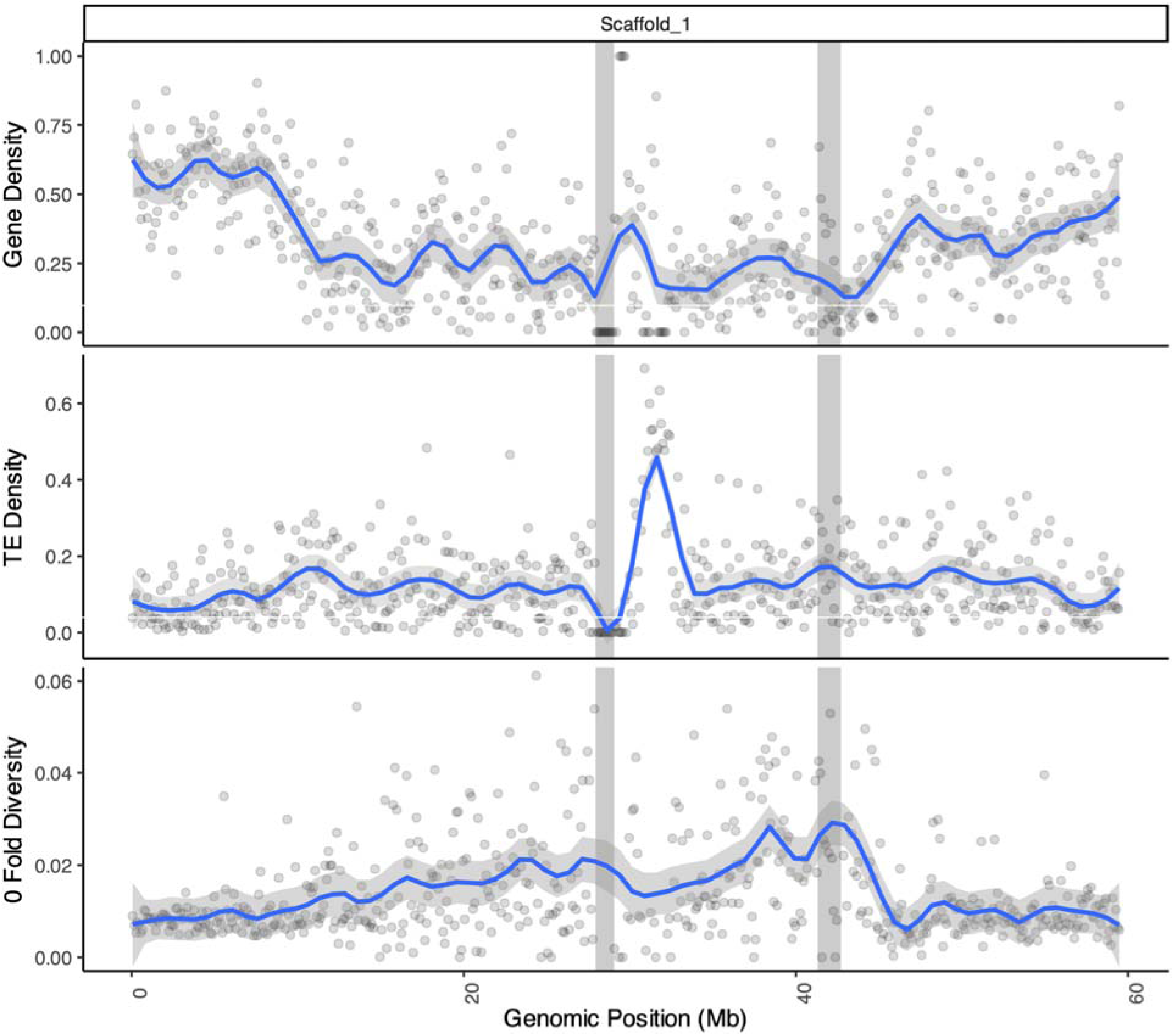
Characterization of chromosome 1, containing the sex determining region between 40-43.5 Mb (right vertical grey bar), a region which shows the highest average 0-fold diversity and among the lowest gene density across chromosome 1. Analysis was done by mapping all reads to only haplotype 2 (i.e. no competitive mapping). Transposon element (TE) density is greatly enriched at ∼32 Mb, just neighboring the inferred location of the centromeric region (left vertical bar; as inferred from RepeatObserver).

**Sup Figure 6.**
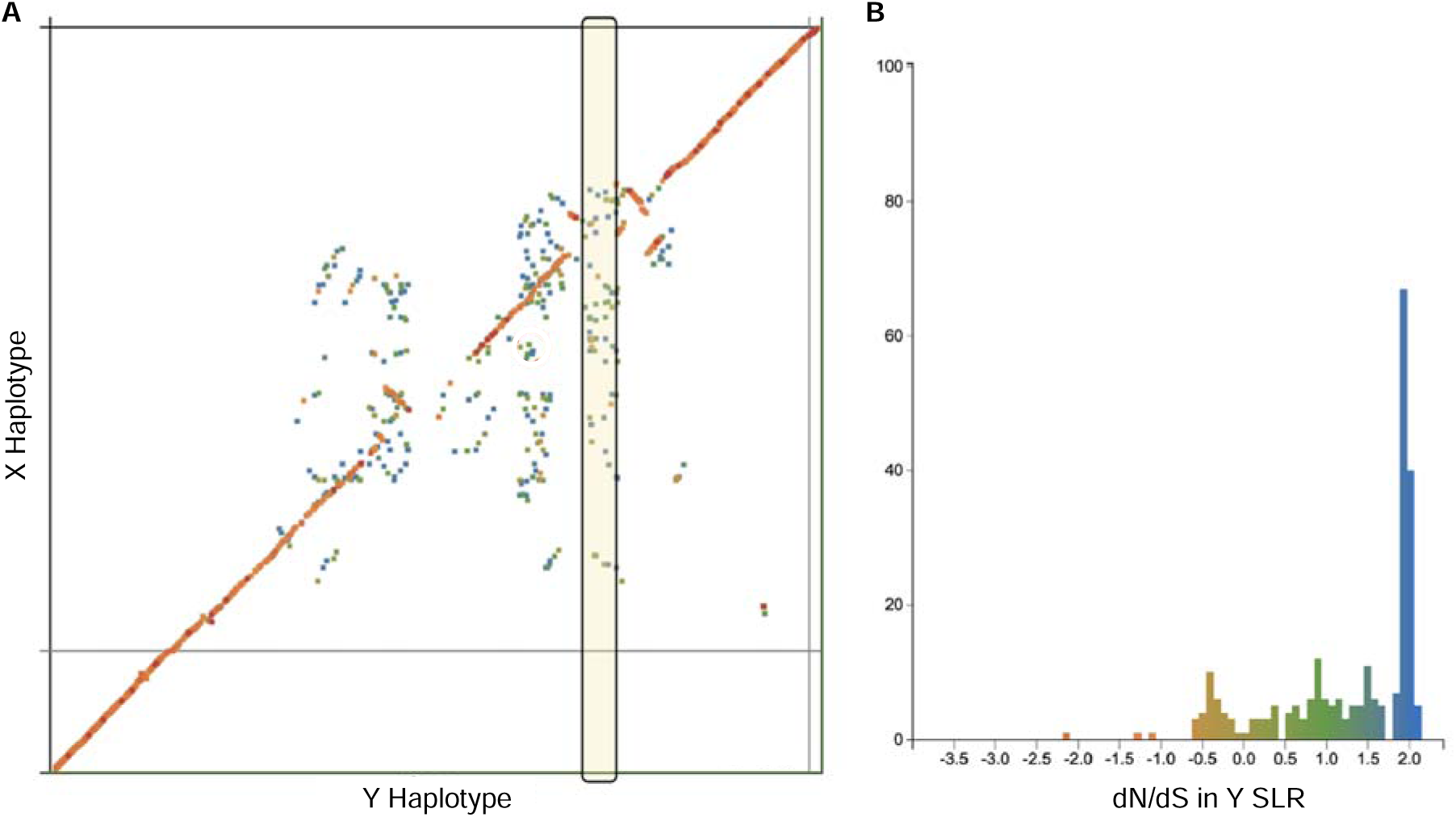
*A)* Genic synteny analyses [92] between the X haplotype and Y of our focal genome resolves the presence of duplicate genes across the X mapping to the Y.) These paralogous show high levels of divergence (dN/dS) consistent with long diverged paralogues.

**Sup Figure 7.**
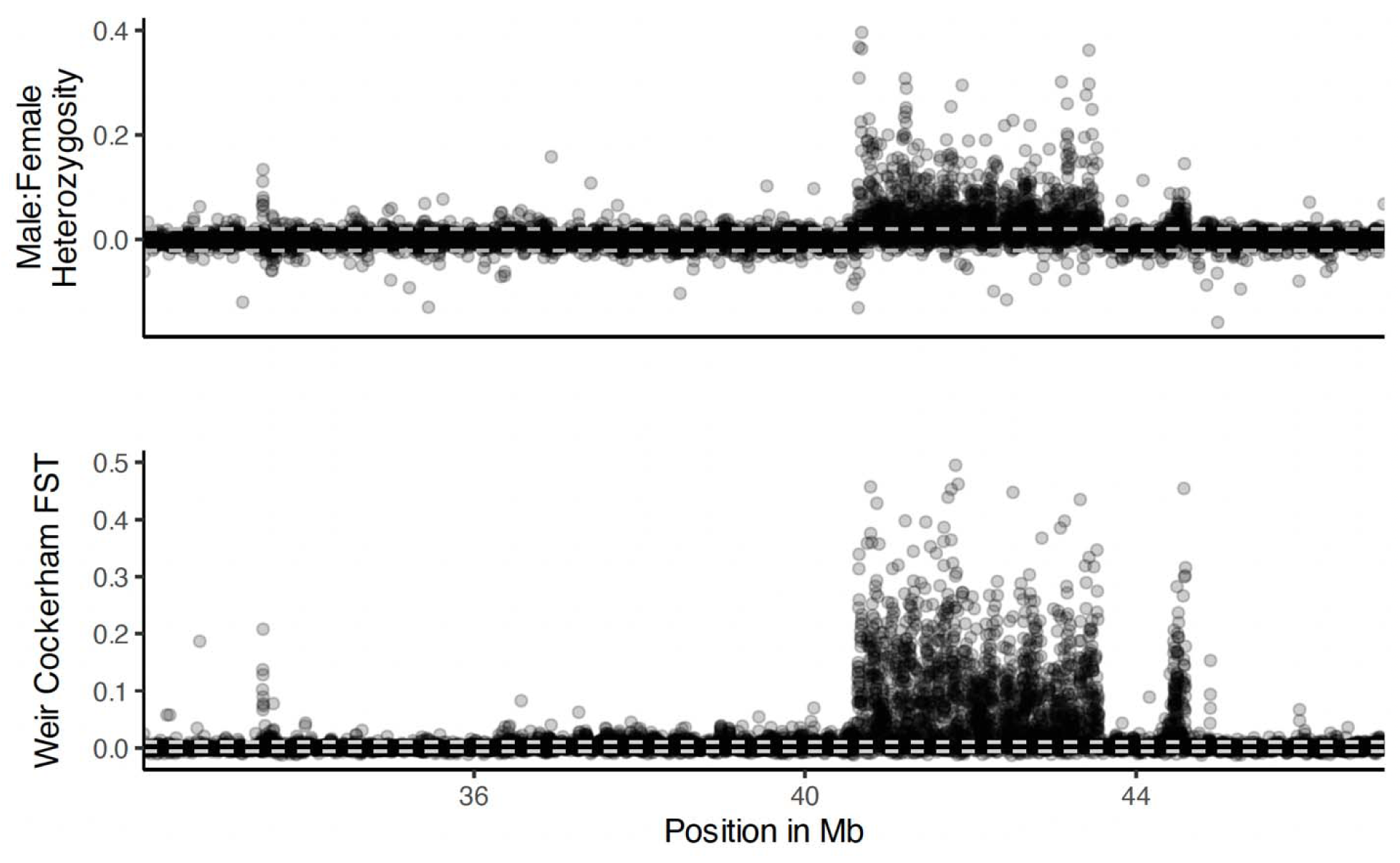
Upper) The difference in heterozygosity between males and females is heightened in the sex-linked region, along with allelic differentiation (F_ST_: Middle Lower) between males and females, where points represent the mean in 1 kb genomic windows. In both plots, the 95% percentile of these summary statistics across autosomes is delimited by the horizontal dashed white lines. Mean F_ST_ in the SLR [40.5-43.5 Mb] is 0.07, whereas the mean ratio of male:female heterozygosity is 0.037.

**Sup Figure 8.**
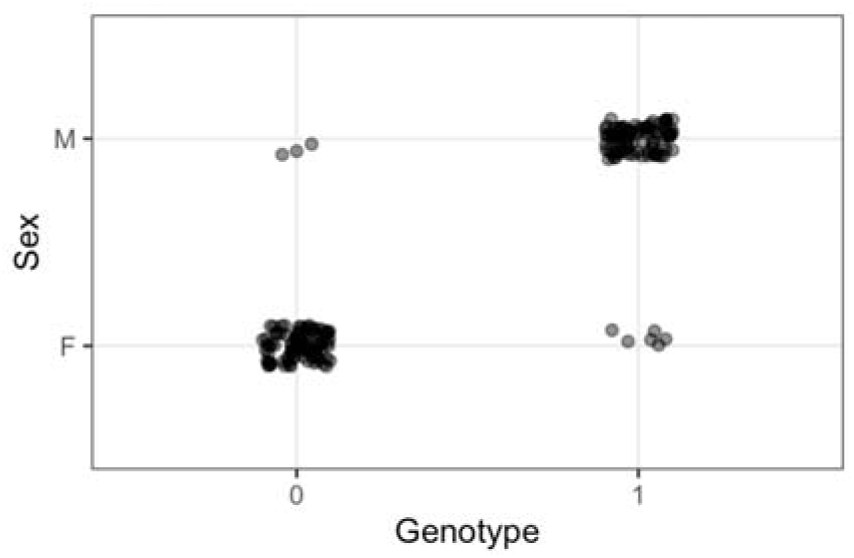
Genotype distribution of the top SNP (genome-wide) in the peak upstream of the primary SLR.

**Sup Figure 9.**
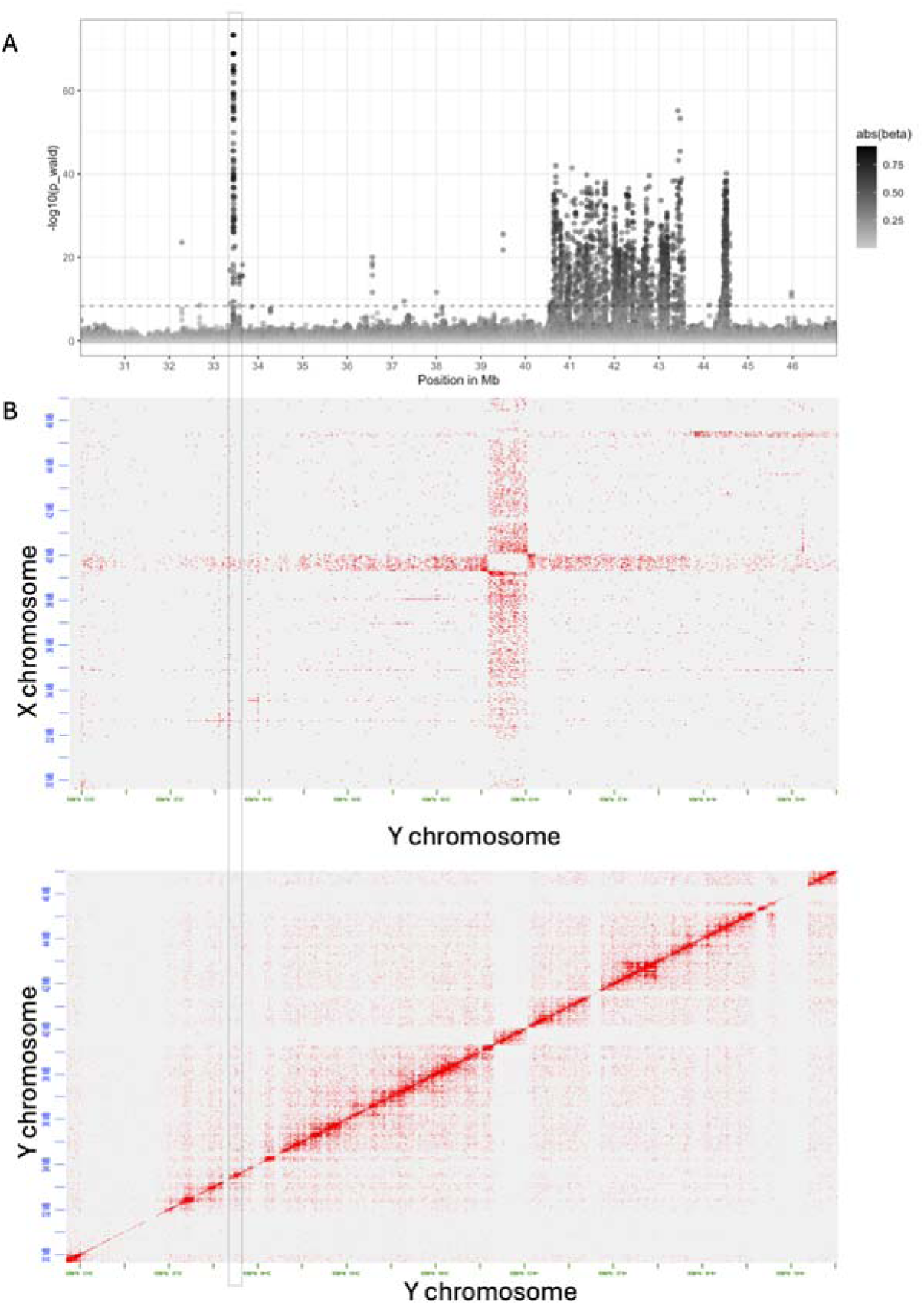
Evaluation of assembly quality near the sex-associated locus on chromosome 1. A) Hi-C contacts between the Y chromosome and the X chromosome support the correct phasing of the SLR, despite an upstream region (not sex-linked) that appears to have been phase switched. B) Contacts within the Y chromosome (lower) support the placement of the sex-associated locus upstream [at ∼33.5Mb] of the primary SLR, and the correct phasing and placement of sex linked contigs within the SLR.

**Sup Figure 10.**
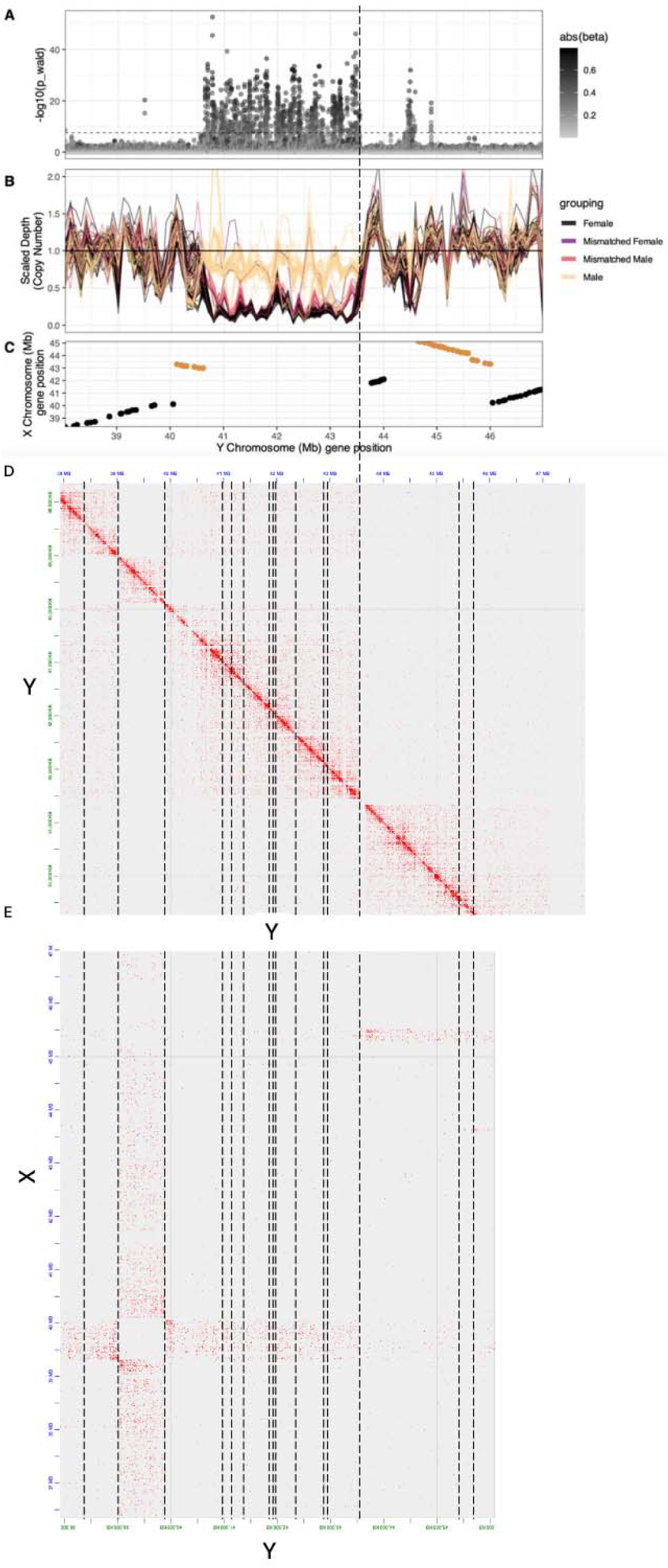
Evaluation of assembly quality near the sex-linked region (SLR) on chromosome 1 (A-C). Hi-C contacts within the Y chromosome (D) support the placement and phasing of the sex-associated peak downstream [at ∼44.5Mb] of the primary SLR. However, the right bound of the SLR does not have contacts with the downstream sequence, suggesting a large gap in our assembly. Contacts between the Y chromosome and the X chromosome (E) show a region with incorrect phasing (highlighted in red) that is not sex-linked. Gaps between scaffolded contigs in the assembly are indicated by vertical dashed lines.

**Sup Figure 11.**
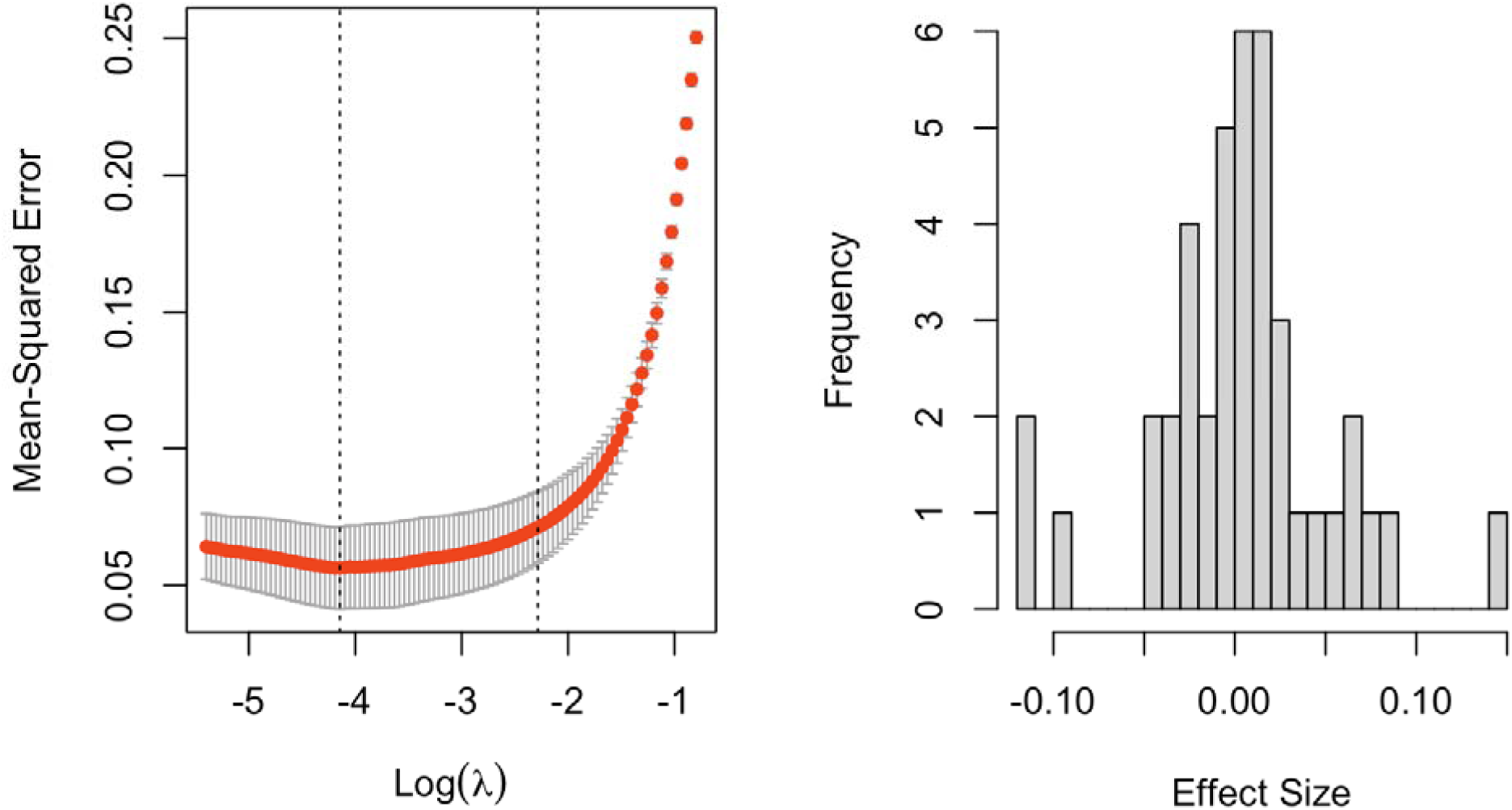
Results for the lasso regression analyses of the predictive independence of loci with significant associations with sex, after Bonferroni correction. Upper) The minimization of MSE occurs when lambda = 0.016. B) The distribution of effect sizes for the 42 loci with remaining non-zero effects on sex.

**Sup Figure 12.**
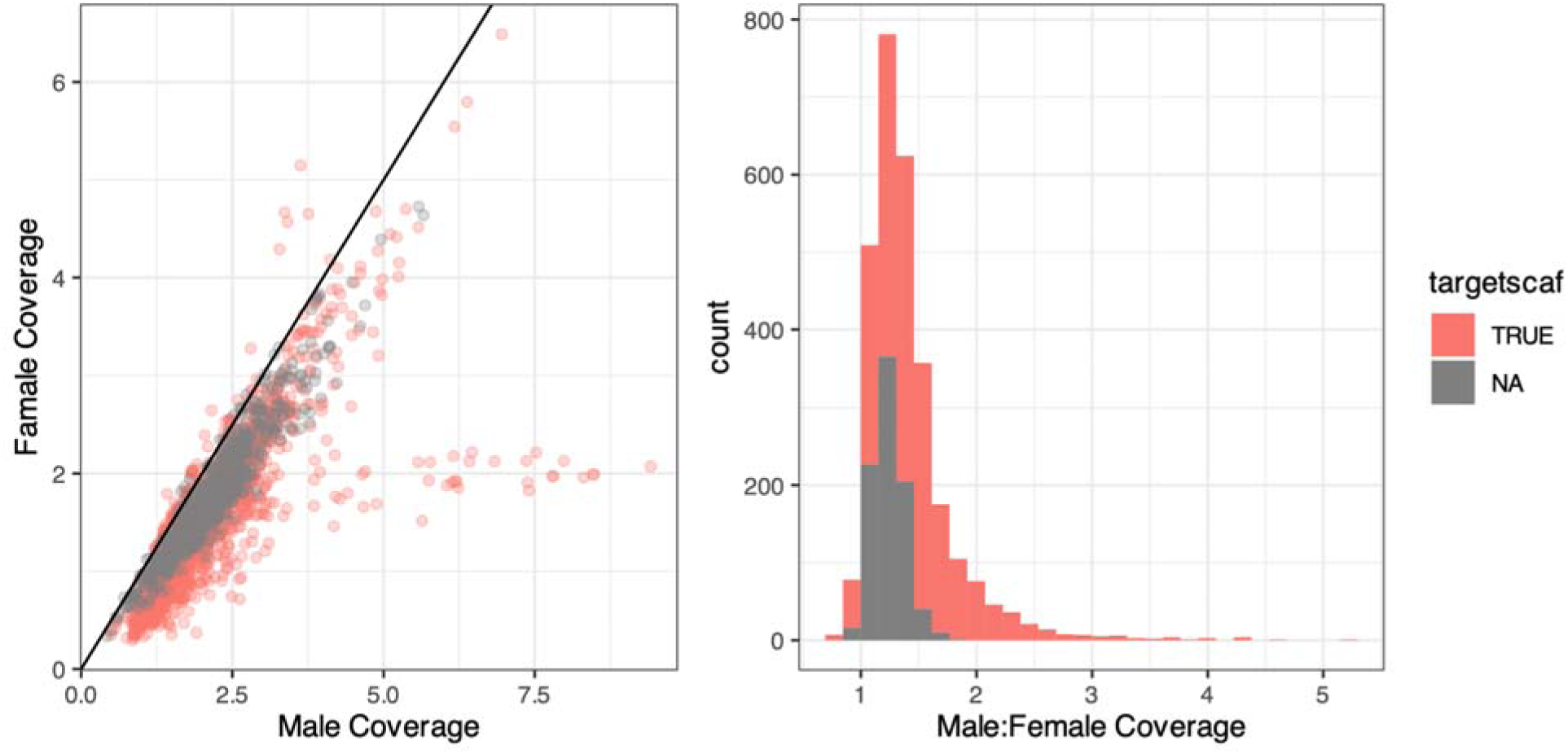
Male versus Female coverage for loci significantly associated with sex (FDR < 0.05). Color indicates whether loci are located on the sex-linked, “target” Chromosome 1, or on other scaffolds across the genome.

**Sup Figure 13.**
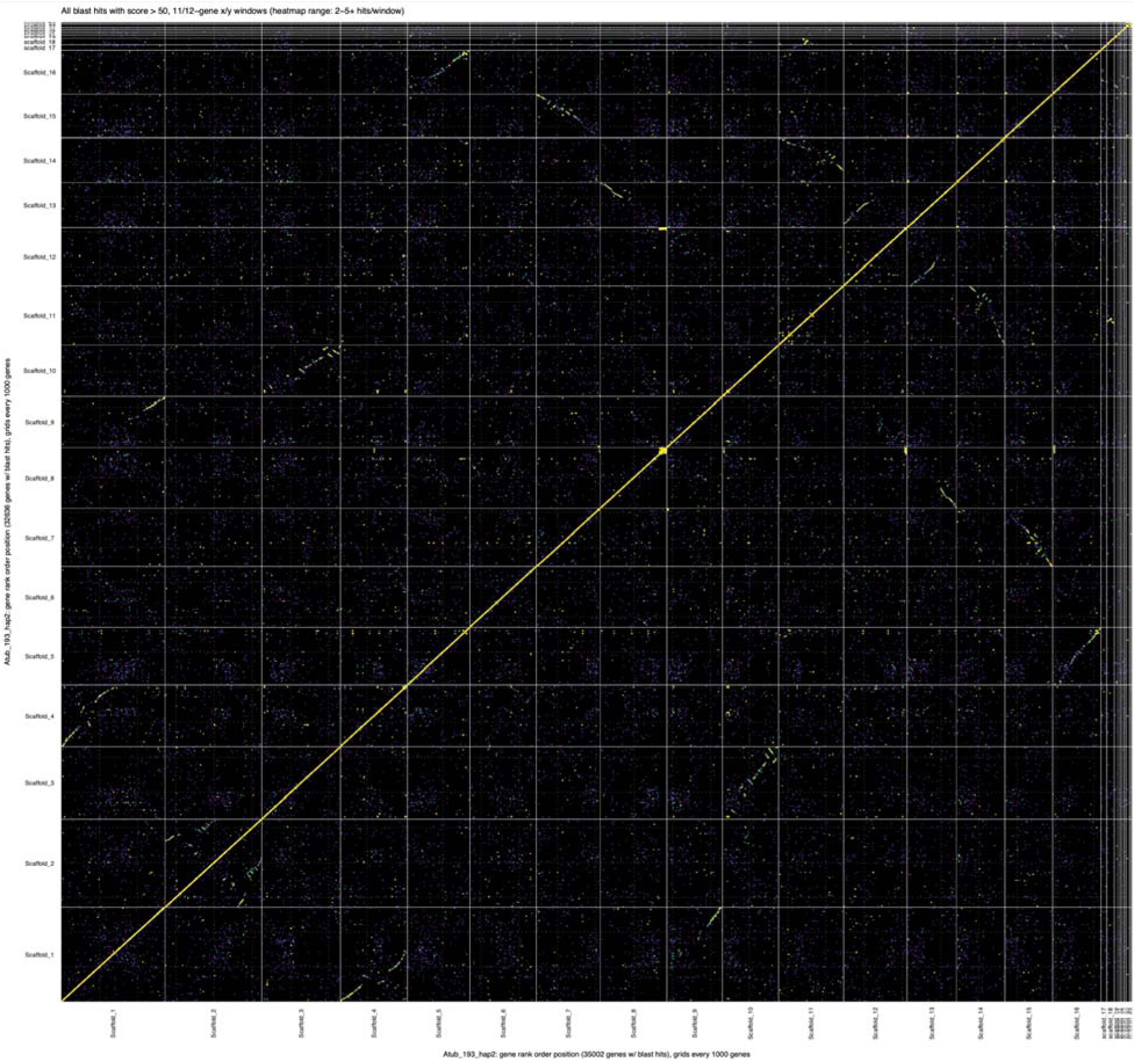
Dotplot of synteny between haplotype 2 and itself (our focal assembly for which population genomic was aligned to, the individual from Walpole]), where dots are gene blast hits with score > 50 inferred from the genespace pipeline. Evidence of a past whole-genome duplication event is evident, as every scaffold shows contiguous stretches of synteny with one or more scaffolds.

**Sup Figure 14.**
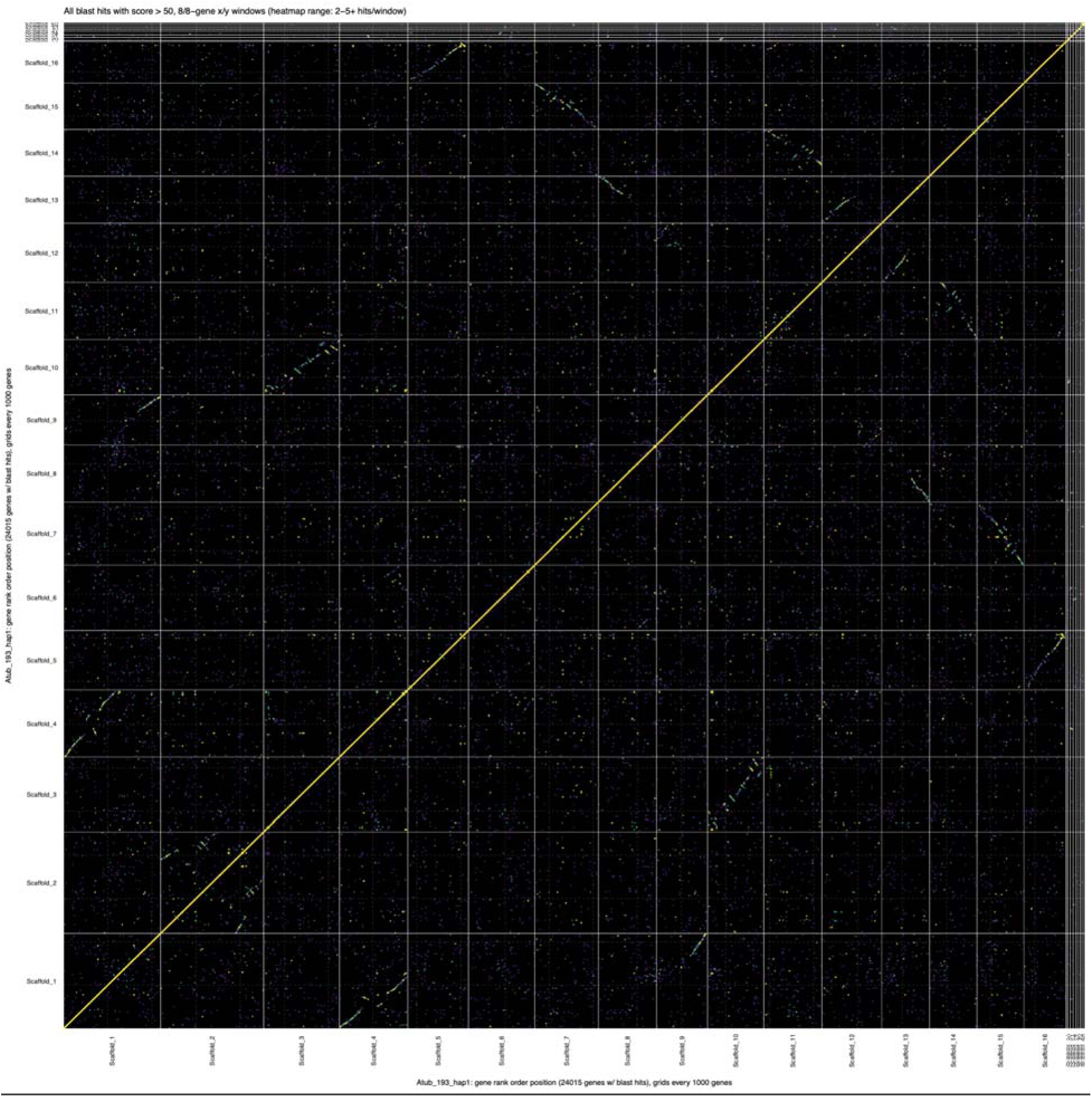
Dotplot of synteny between haplotype 1 against itself [from the admixed Walpole male], where dots are gene blast hits with score > 50 inferred from the genespace pipeline. Evidence of a past whole-genome duplication event is evident, as every scaffold shows contiguous stretches of synteny with one or more scaffolds.

**Sup Figure 15.**
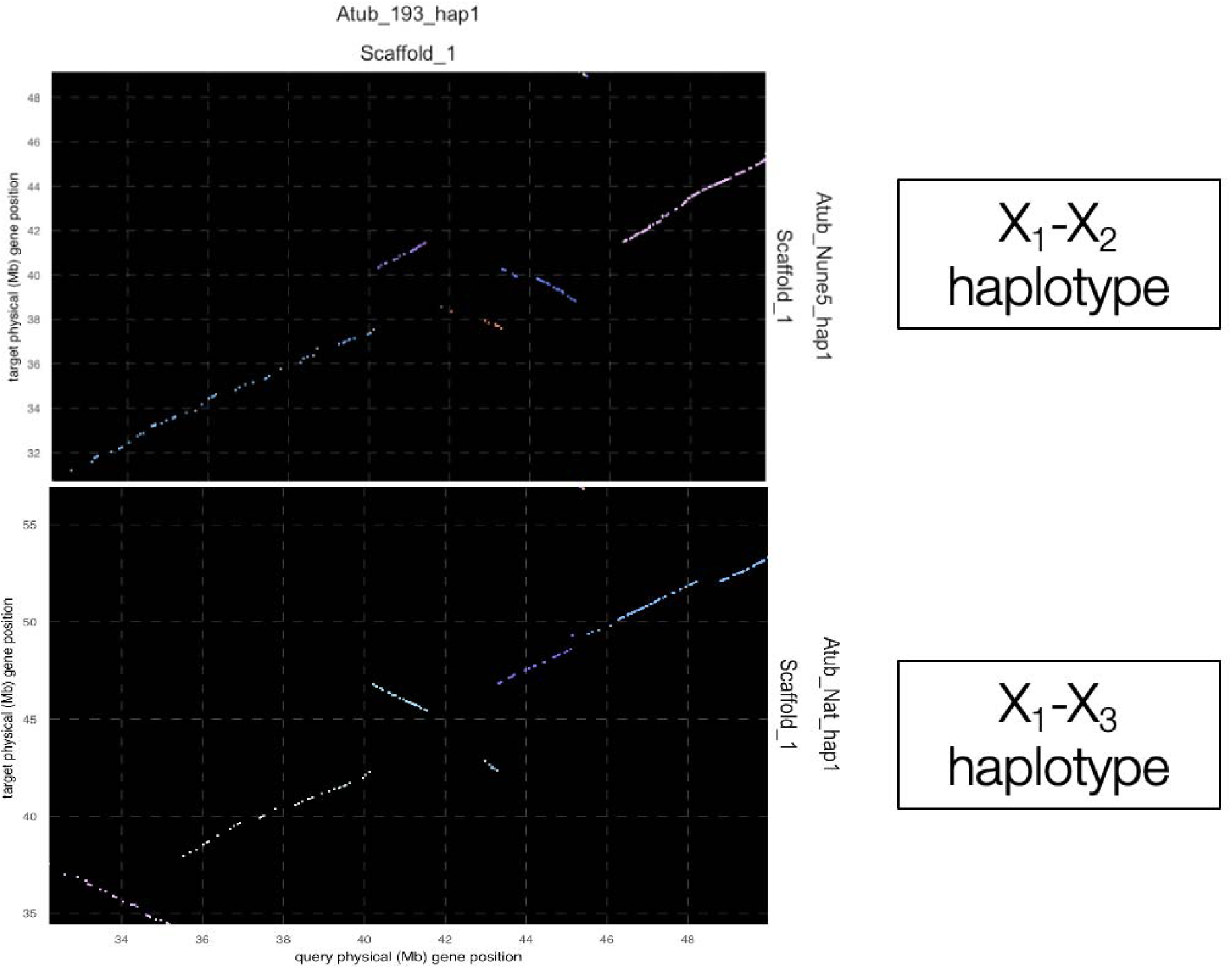
Dotplots of syntenic genes between X haplotype assemblies of the sex-linked region. Colors represent orthologous grouping (i.e. tracts of contiguous syntenic sequence).

**Sup Figure 16.**
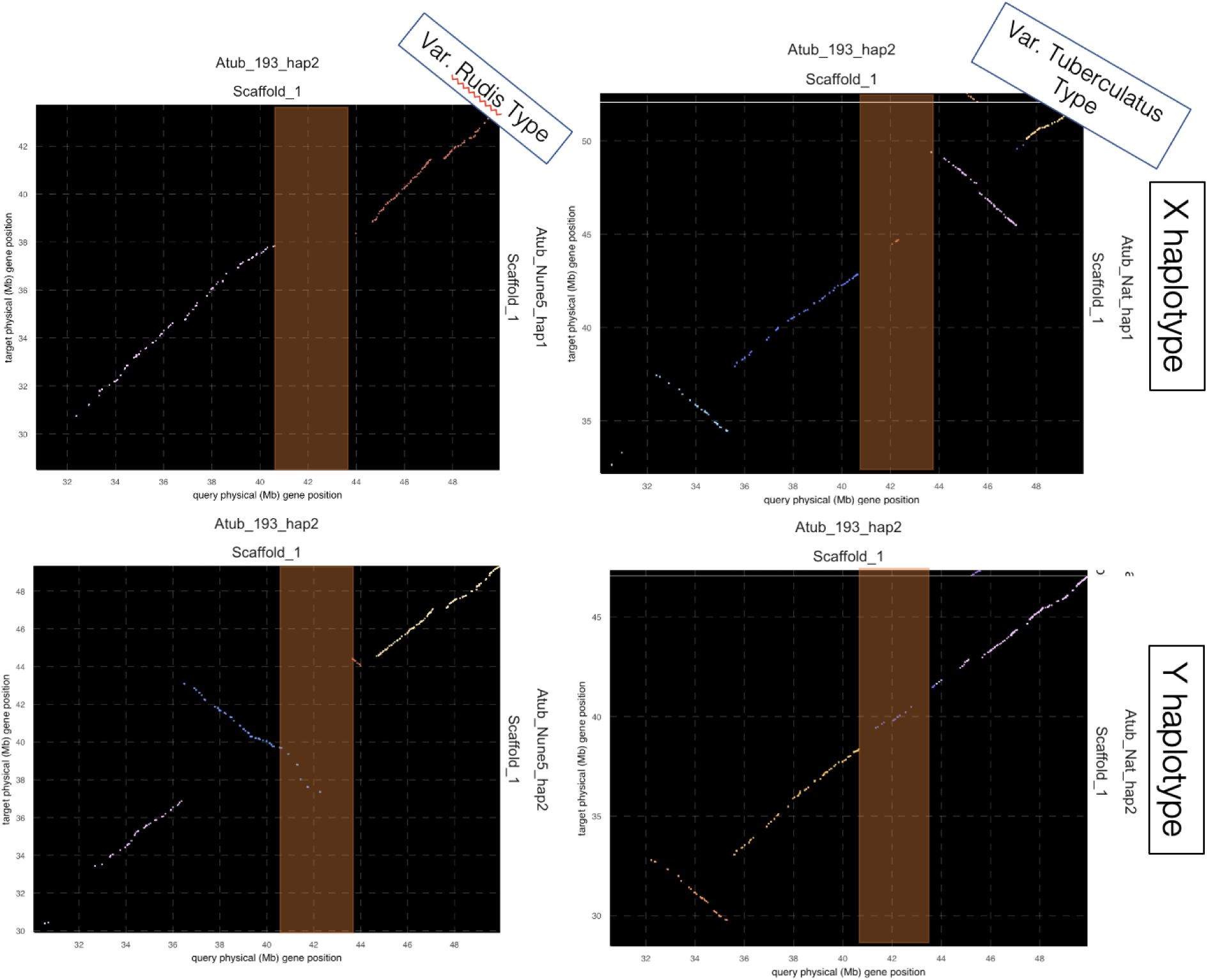
Pairwise comparisons of synteny for our focal-Y (admixed type) and the X and Y haplotypes of the var. *rudis* type individual (left) var. *tuberculatus* type (right).

**Sup Figure 17.**
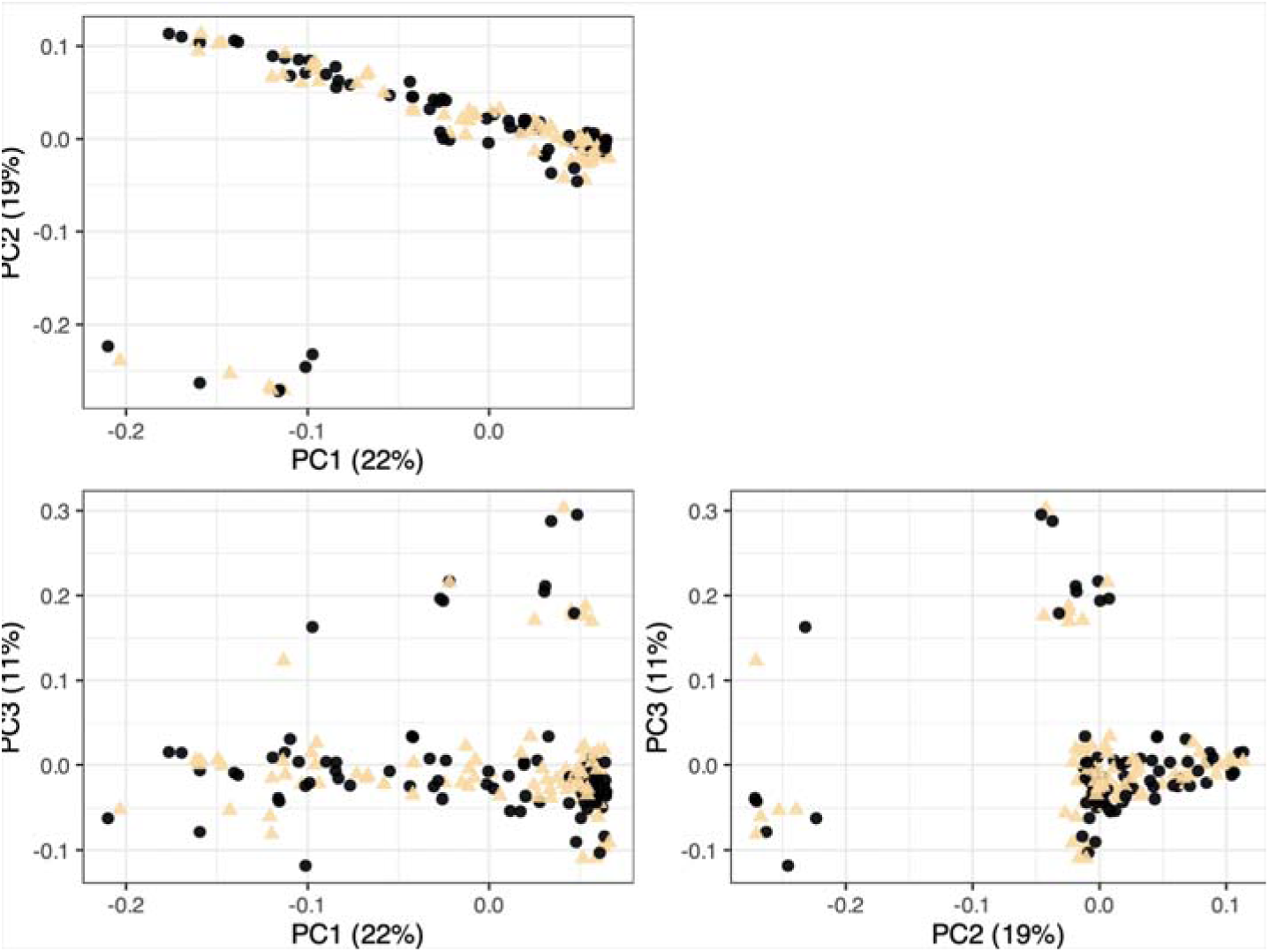
Population structure inferred from genotype information within the inversion (Sup Table 2 number 5* and 14*) just upstream of the sex-linked region. Points (individual resequenced plants) coloured by phenotypic sex assignment (cream = male, black = female), illustrating the lack of sex-based structure in this inversion.

**Sup Figure 18.**
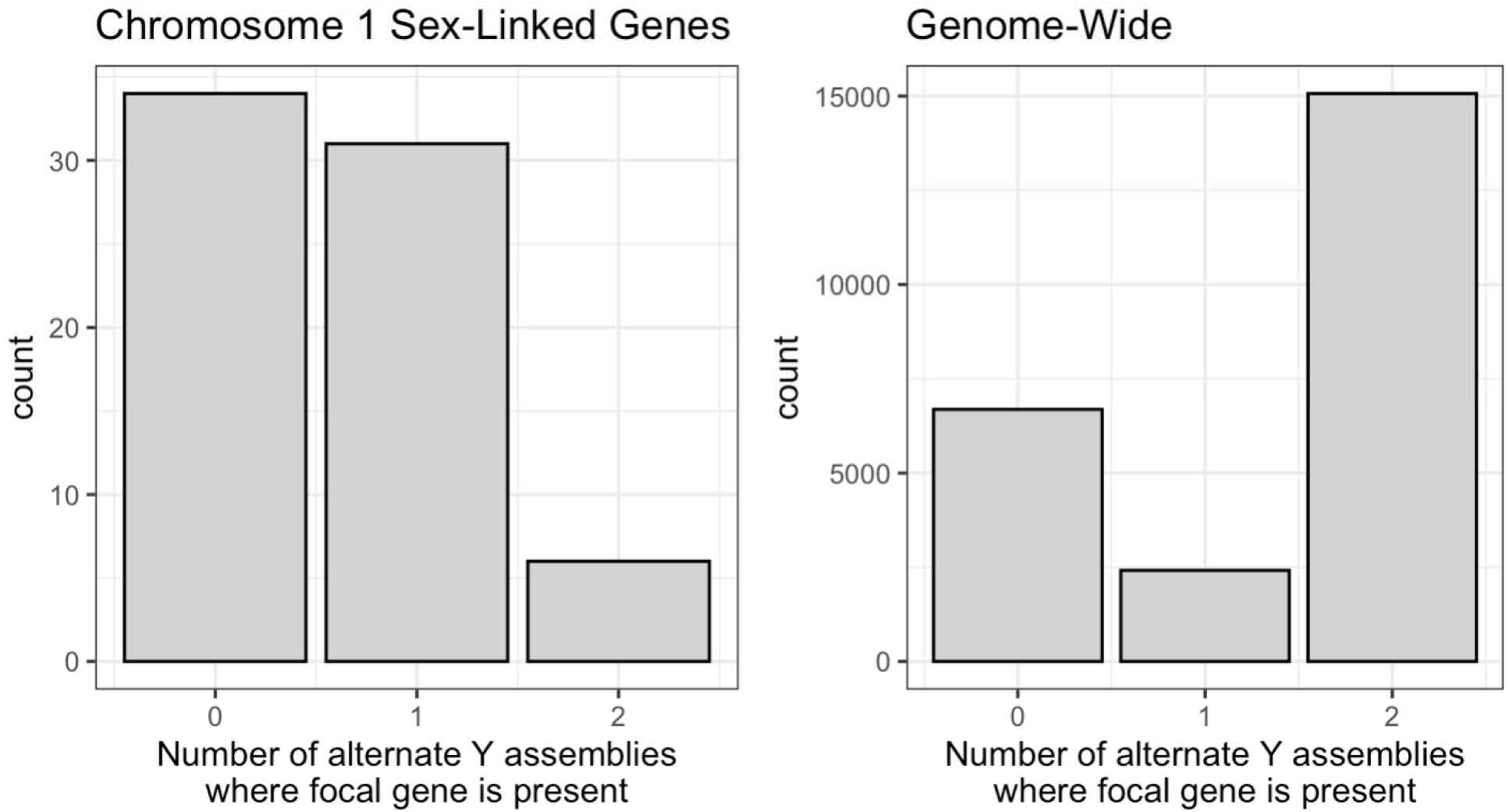
Pairwise comparisons of the focal Y-containing assembly to the two other Y-containing assemblies show enriched gene presence-absence variation in the sex-linked region, compared to genome wide.

**Sup Figure 19.**
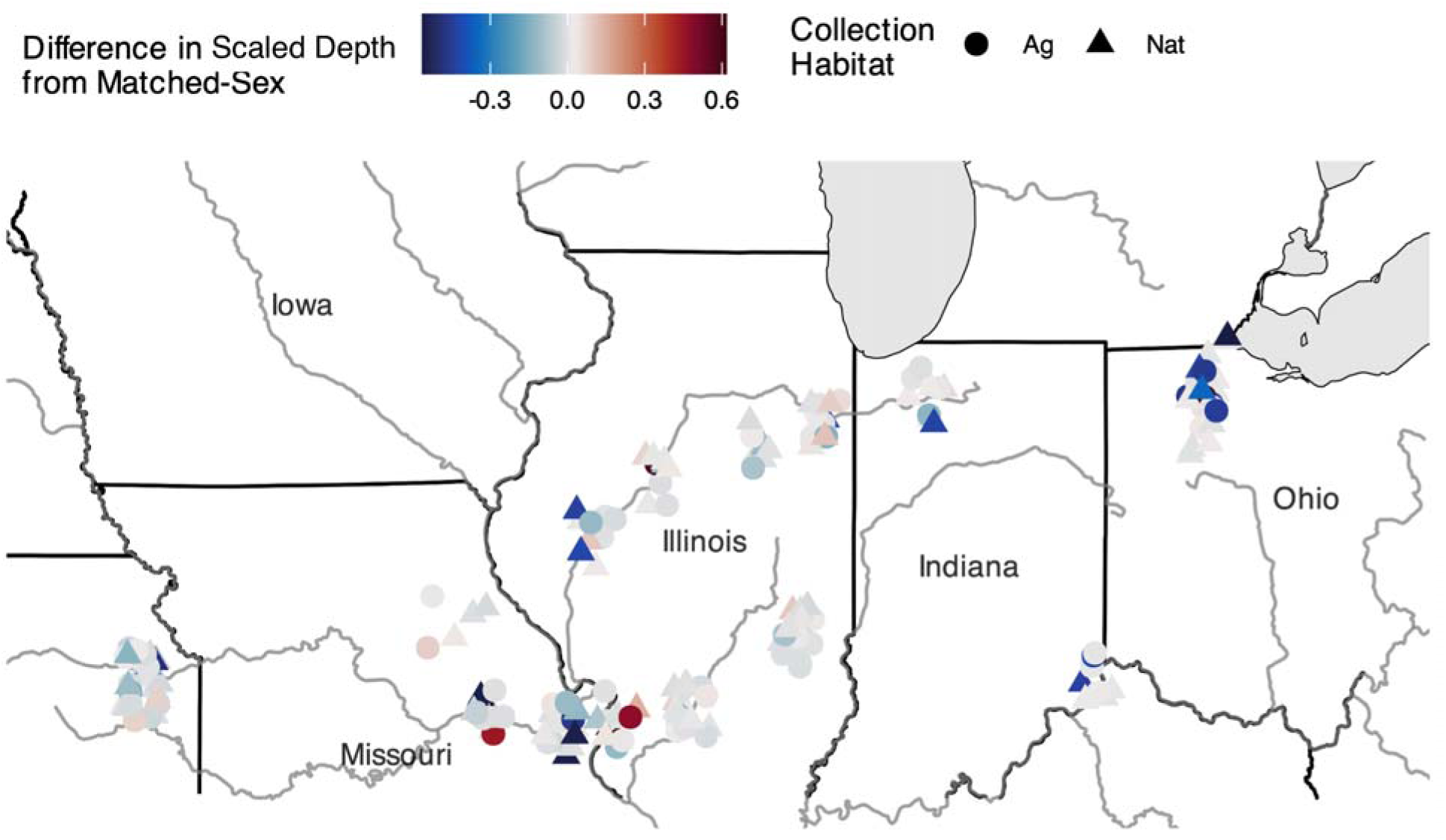
The geographic distribution of the difference in mean normalized read depth across the sex-linked region (SLR; top) between an individual and the mean across individuals of their respective sex-matched genotype. Color coding represents the degree of mismatch based on an individual’s mean normalized read depth across the region, and the mean normalized read depth of the average sex-matched individual. Negative numbers indicate males who genotypically resemble most females, and who therefore have less sequence content in the SLR than most males, whereas positive values indicate females who genotypically resemble most males (have an excess of sequence content in the SLR). Administrative boundaries obtained from GADM (https://gadm.org/) and Natural Earth (https://www.naturalearthdata.com/), while natural features (lakes, rivers, ocean) were obtained from Natural Earth. Both datasets are freely available for academic use and compatible with CC BY 4.0 licensing.

**Sup Figure 20.**
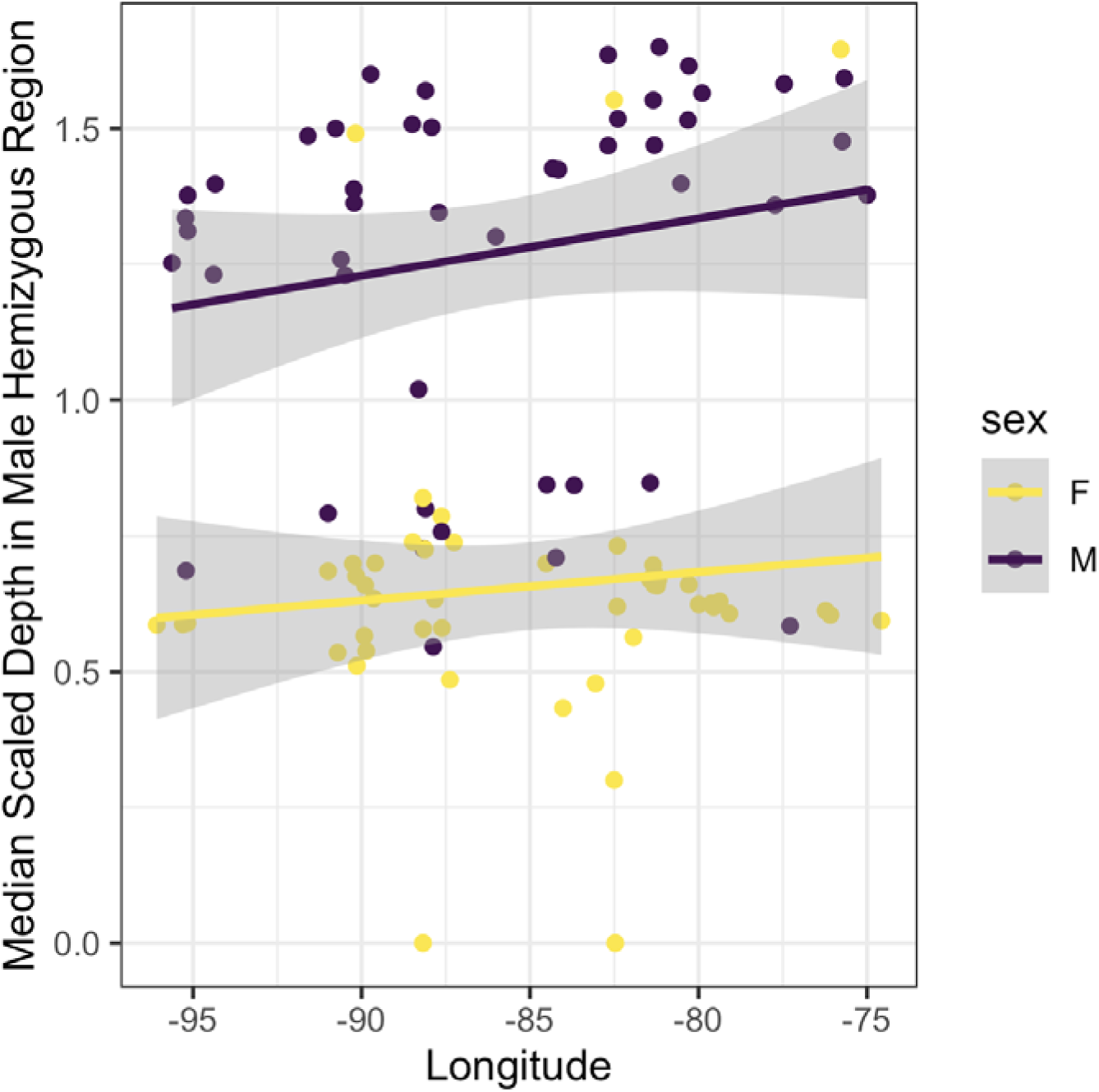
Median scaled depth in sex linked region for an independent dataset [106], illustrating genotype-phenotype mismatch. 10 males show depth profiles more similar to females than expected, and 3 females show depth profiles more similar to females.

**Sup Figure 21.**
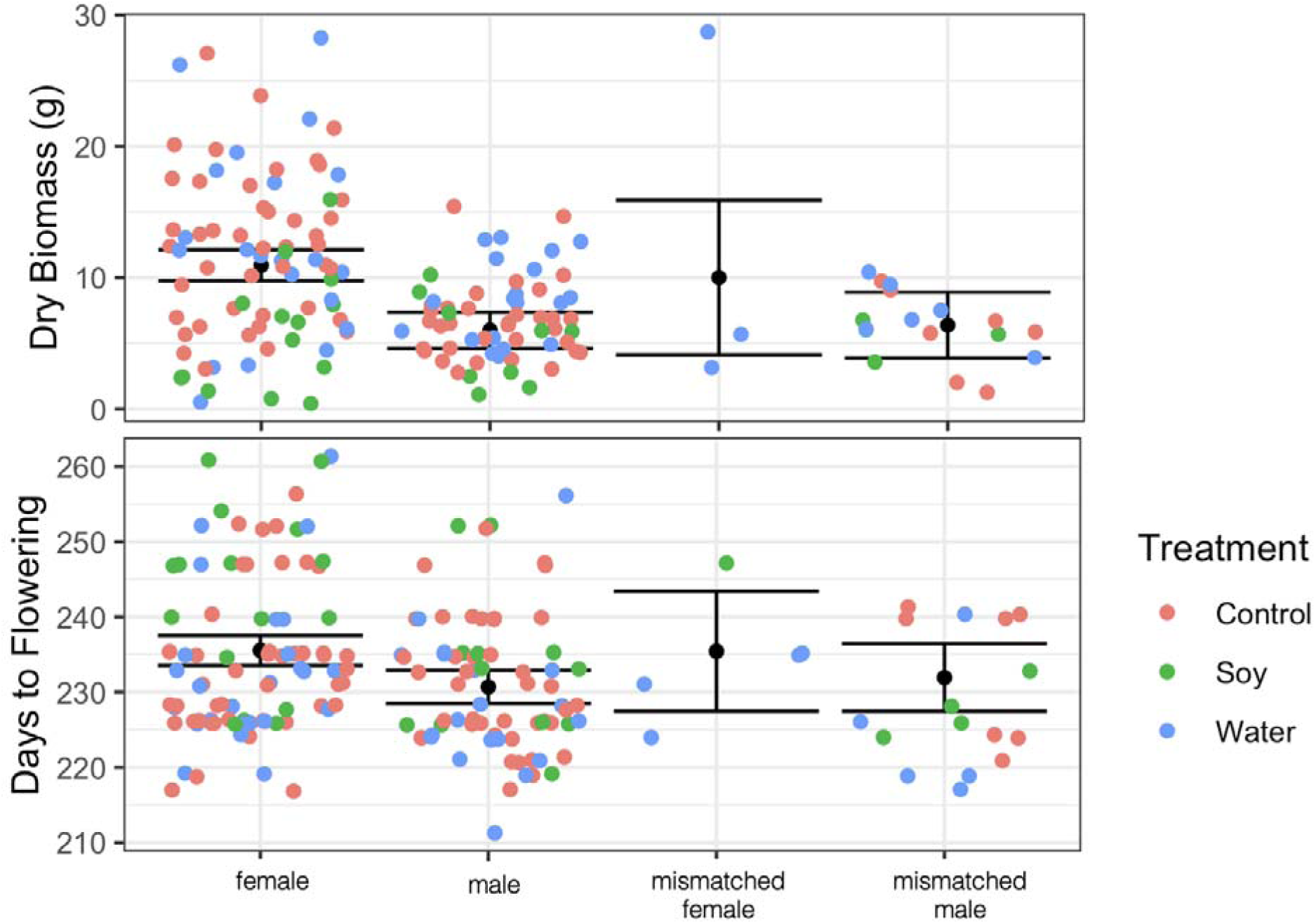
The distribution of biomass and days to flowering for individuals with different phenotype-genotype groupings for sex. We see no significant difference in phenotypes between mismatched individuals and their respective matched sex.

**Sup Figure 22.**
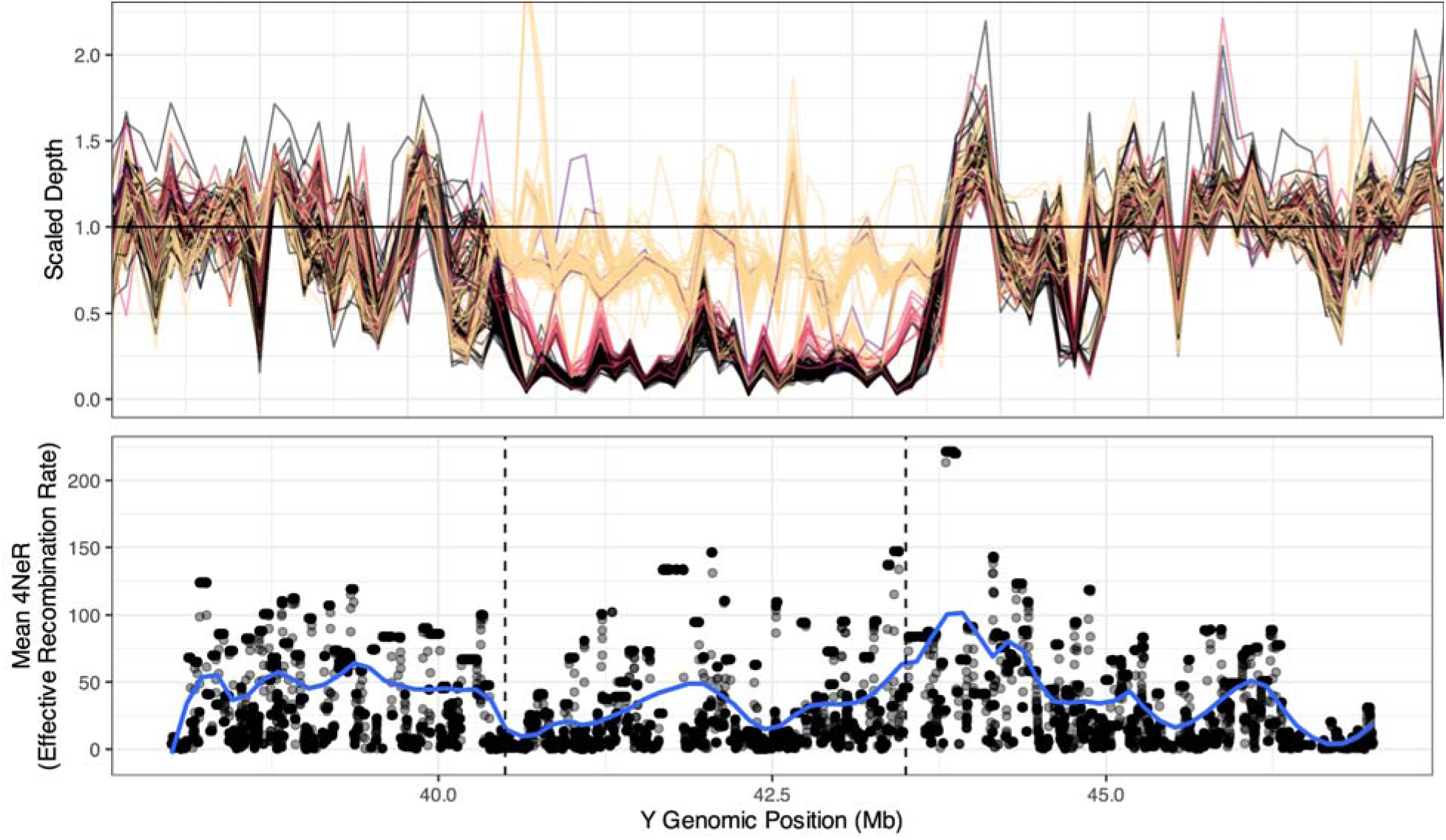
LDhat inferred effective recombination rate as it relates to the SLR. The sex linked region (illustrated by the difference in mapping depth between males and females (top), shows a reduction in effective recombination rate, as estimated between SNPs along the sex linked region on haplotype 2 (the Y containing haplotype) of Scaffold 1.Vertical black dashed lines delimit the SLR.

**Sup Figure 23.**
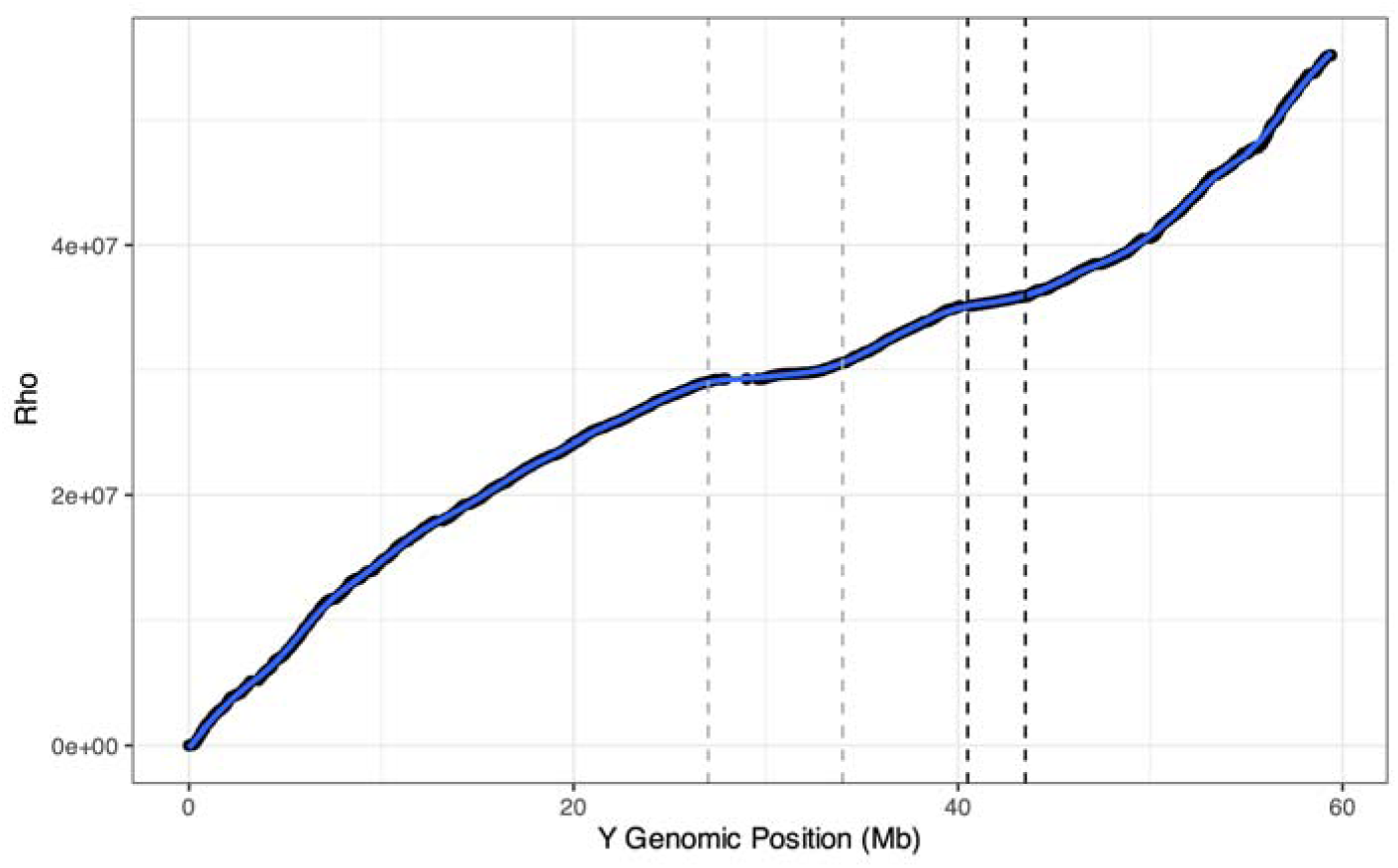
Cumulative rho across the entire proto-Y chromosome (haplotype 2 of Scaffold 1), with points illustrating this value 100 kb windows. Vertical grey dashed lines represent the centromere, whereas vertical black dashed lines represent the SLR.

### Supplementary Tables

**Sup Table 1.**
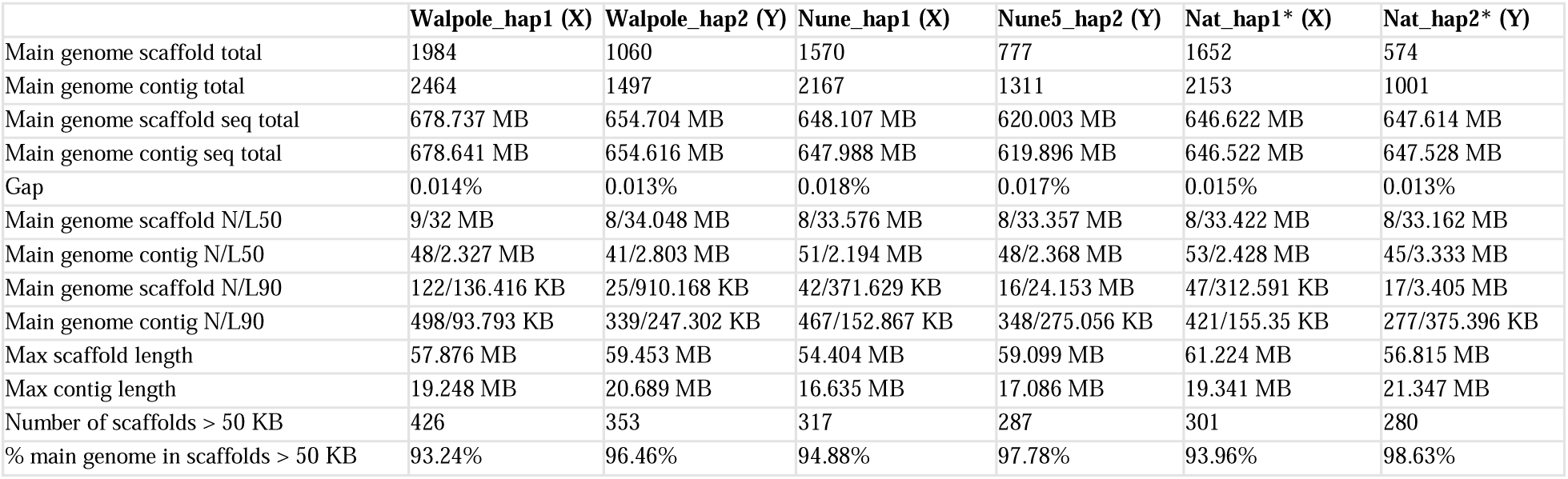
Statistics for 6 haplotype-phased chromosome-level assemblies, from 3 male *A. tuberculatus.* *indicates that the initial HiFiasm assembly step was performed without Hi-C data.

|  | Walpole_hap1 (X) | Walpole_hap2 (Y) | Nune_hap1 (X) | Nune5_hap2 (Y) | Nat_hap1* (X) | Nat_hap2* (Y) |
| --- | --- | --- | --- | --- | --- | --- |
| Main genome scaffold total | 1984 | 1060 | 1570 | 777 | 1652 | 574 |
| Main genome contig total | 2464 | 1497 | 2167 | 1311 | 2153 | 1001 |
| Main genome scaffold seq total | 678.737 MB | 654.704 MB | 648.107 MB | 620.003 MB | 646.622 MB | 647.614 MB |
| Main genome contig seq total | 678.641 MB | 654.616 MB | 647.988 MB | 619.896 MB | 646.522 MB | 647.528 MB |
| Gap | 0.014% | 0.013% | 0.018% | 0.017% | 0.015% | 0.013% |
| Main genome scaffold N/L50 | 9/32 MB | 8/34.048 MB | 8/33.576 MB | 8/33.357 MB | 8/33.422 MB | 8/33.162 MB |
| Main genome contig N/L50 | 48/2.327 MB | 41/2.803 MB | 51/2.194 MB | 48/2.368 MB | 53/2.428 MB | 45/3.333 MB |
| Main genome scaffold N/L90 | 122/136.416 KB | 25/910.168 KB | 42/371.629 KB | 16/24.153 MB | 47/312.591 KB | 17/3.405 MB |
| Main genome contig N/L90 | 498/93.793 KB | 339/247.302 KB | 467/152.867 KB | 348/275.056 KB | 421/155.35 KB | 277/375.396 KB |
| Max scaffold length | 57.876 MB | 59.453 MB | 54.404 MB | 59.099 MB | 61.224 MB | 56.815 MB |
| Max contig length | 19.248 MB | 20.689 MB | 16.635 MB | 17.086 MB | 19.341 MB | 21.347 MB |
| Number of scaffolds > 50 KB | 426 | 353 | 317 | 287 | 301 | 280 |
| % main genome in scaffolds > 50 KB | 93.24% | 96.46% | 94.88% | 97.78% | 93.96% | 98.63% |

**Sup Table 2.** List of Inversions relative to Atub_Walpole_hap2. Note that letter code demarcates the same inversion present across multiple pairwise assembly comparisons (each referenced by the inversion number and alt_genome). Inversion C (5 and 14) represent the variant directly neighboring the SLR in Atub_Walpole_hap2. Predictors of population structure within each inversion were tested in multiple linear regressions of the PC1, PC2, and PC3 of genotypes within the genomic region (significance codes: 0 ‘***’ < 0.001 ‘**’ < 0.01 ‘*’ < 0.05 ‘.’ < 0.1)

| Inversion Number | Ref_Scaf | Start | End | Alt_Genome | Significant Predictors |
| --- | --- | --- | --- | --- | --- |
| 1 | Scaffold_1 | 25970434 | 30554490 | Atub_Walpole_hap1 | NA |
| 2A | Scaffold_1 | 7153812 | 7609088 | Atub_Walpole_hap1 | NA |
| 3 | Scaffold_1 | 40126975 | 40143010 | Atub_Walpole_hap1 | Ancestry<sub>PC2</sub>(.) |
| 4B | Scaffold_1 | 11910786 | 11950873 | Atub_Walpole_hap1 | NA |
| 5C | Scaffold_1 | 44656825 | 46017919 | Atub_Walpole_hap1 | NA |
| 6A | Scaffold_1 | 7153812 | 7609088 | Atub_Nune_hap1 | NA |
| 7 | Scaffold_1 | 36429003 | 43364478 | Atub_Nune_hap1 |  |
| 8 | Scaffold_1 | 43375038 | 44017851 | Atub_Nune_hap1 | Lat<sub>PC2</sub>(.), Env<sub>PC3</sub>(.) |
| 9A | Scaffold_1 | 7153812 | 7609088 | Atub_Nune_hap2 | NA |
| 10A | Scaffold_1 | 7153812 | 7609088 | Atub_Nat_hap1 | NA |
| 11 | Scaffold_1 | 11910786 | 12359692 | Atub_Nat_hap1 | Long<sub>PC1</sub>(*) |
| 12D | Scaffold_1 | 20165508 | 21398392 | Atub_Nat_hap1 | Long<sub>PC1</sub>(*), Anc<sub>PC2</sub>(*), Long<sub>PC3</sub>(.) |
| 13 | Scaffold_1 | 32372322 | 35290168 | Atub_Nat_hap1 | NA |
| 14C | Scaffold_1 | 43649837 | 47205224 | Atub_Nat_hap1 | NA |
| 15A | Scaffold_1 | 7153812 | 7609088 | Atub_Nat_hap2 | NA |
| 16D | Scaffold_1 | 20165508 | 21398392 | Atub_Nat_hap2 | Long<sub>PC1</sub>(*), Anc<sub>PC2</sub>(*), Long<sub>PC3</sub>(.) |
| 17 | Scaffold_1 | 21953637 | 23506917 | Atub_Nat_hap2 | NA |
| 18 | Scaffold_1 | 32218868 | 35290168 | Atub_Nat_hap2 | NA |

## References

1. Dorken ME, Barrett SCH. Sex determination and the evolution of dioecy from monoecy in Sagittaria latifolia (Alismataceae). Proc Biol Sci. 2004;271: 213–219.

2. Lloyd DG. The transmission of genes via pollen and ovules in gynodioecious angiosperms. Theor Popul Biol. 1976;9: 299–316.

3. Charlesworth D. Evolution of recombination rates between sex chromosomes. Philos Trans R Soc Lond B Biol Sci. 2017;372. doi:10.1098/rstb.2016.0456

4. Renner SS, Müller NA. Plant sex chromosomes defy evolutionary models of expanding recombination suppression and genetic degeneration. Nat Plants. 2021;7: 392–402.

5. Ohyama K, Takemura M, Oda K, Fukuzawa H, Kohchi T, Nakayama S, et al. Gene content, organization and molecular evolution of plant organellar genomes and sex chromosomes — Insights from the case of the liverwort Marchantia polymorpha. Proc Jpn Acad Ser B Phys Biol Sci. 2009;85: 108–124.

6. Akagi T, Fujita N, Masuda K, Shirasawa K, Nagaki K, Horiuchi A, et al. Rapid and dynamic evolution of a giant Y chromosome in Silene latifolia bioRxiv. 2023. p. 2023.09.21.558759. doi:10.1101/2023.09.21.558759

7. Filatov DA. Heterochiasmy and Sex Chromosome Evolution in Silene. Genes. 2023;14. doi:10.3390/genes14030543

8. Hough J, Hollister JD, Wang W, Barrett SCH, Wright SI. Genetic degeneration of old and young Y chromosomes in the flowering plant Rumex hastatulus. Proc Natl Acad Sci U S A. 2014;111: 7713–7718.

9. Sacchi B, Humphries Z, Kružlicová J, Bodláková M, Pyne C, Choudhury BI, et al. Phased Assembly of Neo-Sex Chromosomes Reveals Extensive Y Degeneration and Rapid Genome Evolution in Rumex hastatulus. Mol Biol Evol. 2024;41. doi:10.1093/molbev/msae074

10. Harkess A, Zhou J, Xu C, Bowers JE, Van der Hulst R, Ayyampalayam S, et al. The asparagus genome sheds light on the origin and evolution of a young Y chromosome. Nat Commun. 2017;8: 1279.

11. Liu Z, Moore PH, Ma H, Ackerman CM, Ragiba M, Yu Q, et al. A primitive Y chromosome in papaya marks incipient sex chromosome evolution. Nature. 2004;427: 348–352.

12. Charlesworth D, Charlesworth B. Population genetics of partial male-sterility and the evolution of monoecy and dioecy. Heredity. 1978;41: 137–153.

13. Lloyd DG. THE DISTRIBUTIONS OF GENDER IN FOUR ANGIOSPERM SPECIES ILLUSTRATING TWO EVOLUTIONARY PATHWAYS TO DIOECY. Evolution. 1980;34: 123–134.

14. Lesaffre T, Pannell Charles Mullon J. A model for the gradual evolution of dioecy and heterogametic sex determination. doi:10.1101/2023.03.24.534076

15. Lesaffre T, Pannell JR, Mullon C. The joint evolution of separate sexes and sexual dimorphism. bioRxiv. 2024. p. 2024.05.31.596835. doi:10.1101/2024.05.31.596835

16. Grosse-Veldmann B, Weigend M. The geometry of gender: hyper-diversification of sexual systems in Urtica L. (Urticaceae). Cladistics. 2018;34: 131–150.

17. Cronk Q. The distribution of sexual function in the flowering plant: from monoecy to dioecy. Philos Trans R Soc Lond B Biol Sci. 2022;377: 20210486.

18. Chen K-H, Pannell JR. Dioecy in a wind-pollinated herb explained by disruptive selection on sex allocation. Evolutionary Biology. bioRxiv; 2025. Available: https://www.biorxiv.org/content/10.1101/2025.01.07.631798v1.full.pdf

19. Charlesworth B, Charlesworth D. A model for the evolution of dioecy and gynodioecy. Am Nat. 1978;112: 975–997.

20. Korpelainen H. Labile sex expression in plants. Biological Reviews. 1998;73: 157–180.

21. Delph LF. Sexual dimorphism in gender plasticity and its consequences for breeding system evolution. Evol Dev. 2003;5: 34–39.

22. Ehlers BK, Bataillon T. “Inconstant males” and the maintenance of labile sex expression in subdioecious plants. New Phytol. 2007;174: 194–211.

23. Russell JRW, Pannell JR. Sex determination in dioecious Mercurialis annua and its close diploid and polyploid relatives. Heredity. 2015;114: 262–271.

24. Cossard GG, Pannell JR. Enhanced leaky sex expression in response to pollen limitation in the dioecious plant Mercurialis annua. J Evol Biol. 2021;34: 416–422.

25. Cossard GG, Gerchen JF, Li X, Cuenot Y, Pannell JR. The rapid dissolution of dioecy by experimental evolution. Curr Biol. 2021;31: 1277–1283.e5.

26. Käfer J, Méndez M, Mousset S. Labile sex expression in angiosperm species with sex chromosomes. Philos Trans R Soc Lond B Biol Sci. 2022;377: 20210216.

27. Ser JR, Roberts RB, Kocher TD. Multiple interacting loci control sex determination in lake Malawi cichlid fish. Evolution. 2010;64: 486–501.

28. Mosyakin SL, Robertson KR. Amaranthus. Flora of North America Editorial Committee, editor. New York: Oxford University Press; 2003.

29. Stetter MG, Schmid KJ. Analysis of phylogenetic relationships and genome size evolution of the Amaranthus genus using GBS indicates the ancestors of an ancient crop. Mol Phylogenet Evol. 2017;109: 80–92.

30. Waselkov KE, Boleda AS, Olsen KM. A Phylogeny of the Genus Amaranthus (Amaranthaceae) Based on Several Low-Copy Nuclear Loci and Chloroplast Regions. Syst Bot. 2018;43: 439–458.

31. Raiyemo DA, Bobadilla LK, Tranel PJ. Genomic profiling of dioecious Amaranthus species provides novel insights into species relatedness and sex genes. BMC Biol. 2023;21: 37.

32. Murray MJ. The Genetics of Sex Determination in the Family Amaranthaceae. Genetics. 1940;25: 409–431.

33. Grant WF. Cytogenetic studies in Amaranthus: I. Cytological aspects of sex determination in dioecious species. Can J Bot. 1959;37: 413–417.

34. Sauer J. RECENT MIGRATION AND EVOLUTION OF THE DIOECIOUS AMARANTHS. Evolution. 1957;11: 11–31.

35. Trucco F, Jeschke MR, Rayburn AL, Tranel PJ. Amaranthus hybridus can be pollinated frequently by A. tuberculatus under field conditions. Heredity. 2005;94: 64–70.

36. Montgomery JS, Giacomini DA, Weigel D, Tranel PJ. Male-specific Y-chromosomal regions in waterhemp (Amaranthus tuberculatus) and Palmer amaranth (Amaranthus palmeri). New Phytol. 2021;229: 3522–3533.

37. Brown PJ, Upadyayula N, Mahone GS, Tian F, Bradbury PJ, Myles S, et al. Distinct genetic architectures for male and female inflorescence traits of maize. PLoS Genet. 2011;7: e1002383.

38. Li S, Yang D, Zhu Y. Characterization and use of male sterility in hybrid rice breeding. J Integr Plant Biol. 2007;49: 791–804.

39. Neves CJ, Matzrafi M, Thiele M, Lorant A, Mesgaran MB, Stetter MG. Male Linked Genomic Region Determines Sex in Dioecious Amaranthus palmeri. Journal of Heredity. 2020. doi:10.1093/jhered/esaa047

40. Kreiner JM, Caballero A, Wright SI, Stinchcombe JR. Selective ancestral sorting and de novo evolution in the agricultural invasion of Amaranthus tuberculatus. Evolution. 2021. doi:10.1111/evo.14404

41. Kreiner JM, Hnatovska S, Stinchcombe JR, Wright SI. Quantifying the role of genome size and repeat content in adaptive variation and the architecture of flowering time in Amaranthus tuberculatus. PLoS Genet. 2023;19: e1010865.

42. Raiyemo DA, Cutti L, Patterson EL, Llaca V, Fengler K, Montgomery JS, et al. A phased chromosome-level genome assembly provides insights into the evolution of sex chromosomes in Amaranthus tuberculatus. bioRxiv. 2024. p. 2024.05.30.596720. doi:10.1101/2024.05.30.596720

43. Nowak K, Morończyk J, Grzyb M, Szczygieł-Sommer A, Gaj MD. miR172 Regulates WUS during Somatic Embryogenesis in Arabidopsis via AP2. Cells. 2022;11. doi:10.3390/cells11040718

44. Lai Y-S, Zhang X, Zhang W, Shen D, Wang H, Xia Y, et al. The association of changes in DNA methylation with temperature-dependent sex determination in cucumber. J Exp Bot. 2017;68: 2899–2912.

45. Shen C, Zhang Y, Li G, Shi J, Wang D, Zhu W, et al. MADS8 is indispensable for female reproductive development at high ambient temperatures in cereal crops. Plant Cell. 2023;36: 65–84.

46. Kojima S, Takahashi Y, Kobayashi Y, Monna L, Sasaki T, Araki T, et al. Hd3a, a rice ortholog of the Arabidopsis FT gene, promotes transition to flowering downstream of Hd1 under short-day conditions. Plant Cell Physiol. 2002;43: 1096–1105.

47. Elphinstone C, Elphinstone R, Todesco M, Rieseberg L. RepeatOBserver: tandem repeat visualization and centromere detection. bioRxiv. 2023. p. 2023.12.30.573697. doi:10.1101/2023.12.30.573697

48. van der Bijl W, Shu J, Goberdhan VS, Sherin LM, Cortázar-Chinarro M, CorralLLópez A, et al. Deep learning reveals the role of copy number variation in the genetic architecture of a highly polymorphic sexual trait. bioRxiv. 2024. doi:10.1101/2023.09.29.560175

49. Charlesworth B, Charlesworth D. The degeneration of Y chromosomes. Philos Trans R Soc Lond B Biol Sci. 2000;355: 1563–1572.

50. Wilson Sayres MA. Genetic diversity on the sex chromosomes. Genome Biol Evol. 2018;10: 1064–1078.

51. Lightfoot DJ, Jarvis DE, Ramaraj T, Lee R, Jellen EN, Maughan PJ. Single-molecule sequencing and Hi-C-based proximity-guided assembly of amaranth (Amaranthus hypochondriacus) chromosomes provide insights into genome evolution. BMC Biol. 2017;15: 74.

52. Charlesworth B. The Evolution of Sex Chromosomes. Science. 1991;251: 1030–1033.

53. Kirkpatrick M. The Evolution of Genome Structure by Natural and Sexual Selection. J Hered. 2017;108: 3–11.

54. Furman BLS, Metzger DCH, Darolti I, Wright AE, Sandkam BA, Almeida P, et al. Sex Chromosome Evolution: So Many Exceptions to the Rules. Genome Biol Evol. 2020;12: 750–763.

55. Lovell JT, Sreedasyam A, Schranz ME, Wilson M, Carlson JW, Harkess A, et al. GENESPACE tracks regions of interest and gene copy number variation across multiple genomes. Elife. 2022;11. doi:10.7554/eLife.78526

56. Carvalho AB, Kim BY, Uno F. Strong sequencing bias in Nanopore and PacBio prevents assembly of Drosophila melanogaster Y-linked genes. bioRxiv. 2025. p. 2025.02.23.639762. doi:10.1101/2025.02.23.639762

57. Hugouvieux V, Blanc-Mathieu R, Janeau A, Paul M, Lucas J, Xu X, et al. SEPALLATA-driven MADS transcription factor tetramerization is required for inner whorl floral organ development. Plant Cell. 2024;36: 3435–3450.

58. McVean G, Auton A. LDhat 2.1: a package for the population genetic analysis of recombination. Department of Statistics, Oxford, OX1 3TG, UK. 2007. Available: https://pdfs.semanticscholar.org/37c0/71c2bcd115bcce4bb2e4eb89a7dbb861c136.pdf

59. Costea M, Weaver SE, Tardif FJ. The Biology of Invasive Alien Plants in Canada. 3. Amaranthus tuberculatus (Moq.) Sauer var. rudis (Sauer) Costea & Tardif. Can J Plant Sci. 2005;85: 507–522.

60. Josephs EB, Lee YW, Stinchcombe JR, Wright SI. Association mapping reveals the role of purifying selection in the maintenance of genomic variation in gene expression. Proc Natl Acad Sci U S A. 2015;112: 15390–15395.

61. Janousek B, Siroký J, Vyskot B. Epigenetic control of sexual phenotype in a dioecious plant, Melandrium album. Mol Gen Genet. 1996;250: 483–490.

62. Feil J. Reproductive ecology of dioecious Siparuna (Monimiaceae) in Ecuador: a case of gall midge pollination. Bot J Linn Soc. 1992;110: 171–203.

63. Renner, S. S., and G. Hausner. Siparunaceae and Monimiacea. In: Andersson GHA, editor. Flora of Ecuador. Sweden: Berlings Arlov; 1997. pp. 1–125.

64. Renner SS, Won H. Repeated evolution of dioecy from monoecy in Siparunaceae (Laurales). Syst Biol. 2001;50: 700–712.

65. Villamil N, Li X, Seddon E, Pannell JR. Simulated herbivory enhances leaky sex expression in the dioecious herb Mercurialis annua. Ann Bot. 2022;129: 79–86.

66. Montgomery JS, Sadeque A, Giacomini DA, Brown PJ, Tranel PJ. Sex-specific markers for waterhemp (Amaranthus tuberculatus) and Palmer amaranth (Amaranthus palmeri). Weed Sci. 2019;67: 412–418.

67. Wu W, Jernstedt J, Mesgaran MB. Comparative floral development in male and female plants of Palmer amaranth (Amaranthus palmeri). Am J Bot. 2023;110: e16212.

68. Groh JS, Vik DC, Stevens KA, Brown PJ, Langley CH, Coop G. Distinct ancient structural polymorphisms control heterodichogamy in walnuts and hickories. bioRxiv. 2024. p. 2023.12.23.573205. doi:10.1101/2023.12.23.573205

69. Blaser O, Grossen C, Neuenschwander S, Perrin N. Sex-chromosome turnovers induced by deleterious mutation load. Evolution. 2013;67: 635–645.

70. Guerrero RF, Rousset F, Kirkpatrick M. Coalescent patterns for chromosomal inversions in divergent populations. Philos Trans R Soc Lond B Biol Sci. 2012;367: 430–438.

71. Kapun M, Mitchell ED, Kawecki TJ, Schmidt P, Flatt T. An Ancestral Balanced Inversion Polymorphism Confers Global Adaptation. Mol Biol Evol. 2023;40. doi:10.1093/molbev/msad118

72. De-Kayne R, Gordon IJ, Terblanche RF, Collins S, Omufwoko KS, Martins DJ, et al. Extensive haplotype diversity in a butterfly colour pattern supergene is fuelled by incomplete recombination suppression. bioRxiv. 2024. p. 2024.07.26.605145. doi:10.1101/2024.07.26.605145

73. Chang C-H, Gregory LE, Gordon KE, Meiklejohn CD, Larracuente AM. Unique structure and positive selection promote the rapid divergence of Drosophila Y chromosomes. Elife. 2022;11. doi:10.7554/eLife.75795

74. Rozen S, Skaletsky H, Marszalek JD, Minx PJ, Cordum HS, Waterston RH, et al. Abundant gene conversion between arms of palindromes in human and ape Y chromosomes. Nature. 2003;423: 873–876.

75. Almeida P, Sandkam BA, Morris J, Darolti I, Breden F, Mank JE. Divergence and Remarkable Diversity of the Y Chromosome in Guppies. Mol Biol Evol. 2021;38: 619–633.

76. Charlesworth B, Barton NH. The Spread of an Inversion with Migration and Selection. Genetics. 2018;208: 377–382.

77. Kapun M, Flatt T. The adaptive significance of chromosomal inversion polymorphisms in Drosophila melanogaster. Mol Ecol. 2019;28: 1263–1282.

78. Charlesworth B, Flatt T. On the fixation or non-fixation of adaptive inversions. Authorea Preprints. Authorea, Inc.; 2021. doi:10.22541/au.161474567.72404976/v1

79. Zhang X, Pan L, Guo W, Li Y, Wang W. A convergent mechanism of sex determination in dioecious plants: Distinct sex-determining genes display converged regulation on floral B-class genes. Front Plant Sci. 2022;13: 953445.

80. Shore P, Sharrocks AD. The MADS-box family of transcription factors. Eur J Biochem. 1995;229: 1–13.

81. Masiero S, Colombo L, Grini PE, Schnittger A, Kater MM. The emerging importance of type I MADS box transcription factors for plant reproduction. Plant Cell. 2011;23: 865–872.

82. Airoldi CA, Davies B. Gene duplication and the evolution of plant MADS-box transcription factors. J Genet Genomics. 2012;39: 157–165.

83. Qiu Y, Li Z, Köhler C. Ancestral duplication of MADS-box genes in land plants empowered the functional divergence between sporophytes and gametophytes. New Phytol. 2024. doi:10.1111/nph.20065

84. Cronk Q, Müller NA. Default Sex and Single Gene Sex Determination in Dioecious Plants. Front Plant Sci. 2020;11: 1162.

85. Kumam Y, Trick HN, Sharma V, Prasad PVV, Jugulam M. Establishment of first protocol of hypocotyl-based regeneration and callus transformation in waterhemp (Amaranthus tuberculatus). In Vitro Cellular & Developmental Biology - Plant. 2024. doi:10.1007/s11627-023-10408-7

86. Kreiner JM, Giacomini DA, Bemm F, Waithaka B, Regalado J, Lanz C, et al. Multiple modes of convergent adaptation in the spread of glyphosate-resistant Amaranthus tuberculatus. Proc Natl Acad Sci U S A. 2019. doi:10.1073/pnas.1900870116

87. Hirabayashi K, Dumigan CR, Kučka M, Percy DM, Guerriero G, Cronk Q, et al. A High-Quality Phased Genome Assembly of Stinging Nettle (Urtica dioica ssp. dioica). Plants. 2025;14: 124.

88. Todesco M, Owens GL, Bercovich N, Légaré J-S, Soudi S, Burge DO, et al. Massive haplotypes underlie ecotypic differentiation in sunflowers. Nature. 2020;584: 602–607.

89. Stoffel K, van Leeuwen H, Kozik A, Caldwell D, Ashrafi H, Cui X, et al. Development and application of a 6.5 million feature Affymetrix Genechip for massively parallel discovery of single position polymorphisms in lettuce (Lactuca spp.). BMC Genomics. 2012;13: 185.

90. Cheng H, Concepcion GT, Feng X, Zhang H, Li H. Haplotype-resolved de novo assembly using phased assembly graphs with hifiasm. Nat Methods. 2021;18: 170–175.

91. Zhou C, McCarthy SA, Durbin R. YaHS: yet another Hi-C scaffolding tool. Bioinformatics. 2023;39. doi:10.1093/bioinformatics/btac808

92. Haug-Baltzell A, Stephens SA, Davey S, Scheidegger CE, Lyons E. SynMap2 and SynMap3D: web-based whole-genome synteny browsers. Bioinformatics. 2017;33: 2197–2198.

93. Cantarel BL, Korf I, Robb SMC, Parra G, Ross E, Moore B, et al. MAKER: an easy-to-use annotation pipeline designed for emerging model organism genomes. Genome Res. 2008;18: 188–196.

94. Adhikary D, Deyholos MK, Délano-Frier JP. The Amaranth Genome. Springer Nature; 2021.

95. Ma X, Vaistij FE, Li Y, Jansen van Rensburg WS, Harvey S, Bairu MW, et al. A chromosome-level Amaranthus cruentus genome assembly highlights gene family evolution and biosynthetic gene clusters that may underpin the nutritional value of this traditional crop. Plant J. 2021;107: 613–628.

96. Wang H, Xu D, Wang S, Wang A, Lei L, Jiang F, et al. Chromosome-scale Amaranthus tricolor genome provides insights into the evolution of the genus Amaranthus and the mechanism of betalain biosynthesis. DNA Res. 2023;30. doi:10.1093/dnares/dsac050

97. Montgomery JS, Giacomini D, Waithaka B, Lanz C, Murphy BP, Campe R, et al. Draft Genomes of Amaranthus tuberculatus, Amaranthus hybridus, and Amaranthus palmeri. Genome Biol Evol. 2020;12: 1988–1993.

98. Li H, Durbin R. Fast and accurate short read alignment with Burrows-Wheeler transform. Bioinformatics. 2009;25: 1754–1760.

99. Danecek P, Bonfield JK, Liddle J, Marshall J. Twelve years of SAMtools and BCFtools. 2021. Available: https://academic.oup.com/gigascience/article-abstract/10/2/giab008/6137722

100. Zhou X, Stephens M. Genome-wide efficient mixed-model analysis for association studies. Nat Genet. 2012;44: 821–824.

101. Korunes KL, Samuk K. pixy: Unbiased estimation of nucleotide diversity and divergence in the presence of missing data. Mol Ecol Resour. 2021;21: 1359–1368.

102. Minh BQ, Schmidt HA, Chernomor O, Schrempf D, Woodhams MD, von Haeseler A, et al. IQ-TREE 2: New Models and Efficient Methods for Phylogenetic Inference in the Genomic Era. Mol Biol Evol. 2020;37: 1530–1534.

103. Wickham H. ggplot2: Elegant Graphics for Data Analysis. Springer Science & Business Media; 2009.

104. Wilke CO. cowplot: Streamlined Plot Theme and Plot Annotations for ggplot2 (2020). R package version 1.1. 1. 2021.

105. Pedersen BS, Quinlan AR. Mosdepth: quick coverage calculation for genomes and exomes. Bioinformatics. 2018;34: 867–868.

106. Kreiner JM, Latorre SM, Burbano HA, Stinchcombe JR, Otto SP, Weigel D, et al. Rapid weed adaptation and range expansion in response to agriculture over the past two centuries. Science. 2022;378: 1079–1085.

107. Peltzer A. Computational methods for ancient genome reconstruction. Eberhard Karls Universität Tübingen. 2019. Available: https://pure.mpg.de/pubman/faces/ViewItemOverviewPage.jsp?itemId=item_3022008

108. Jónsson H, Ginolhac A, Schubert M, Johnson PLF, Orlando L. mapDamage2.0: fast approximate Bayesian estimates of ancient DNA damage parameters. Bioinformatics. 2013;29: 1682–1684.

109. Kahle D, Wickham H. Ggmap: Spatial visualization with ggplot2. R J. 2013;5: 144.

